# Hippocampal volume across age: Nomograms derived from over 19,700 people in UK Biobank

**DOI:** 10.1101/562678

**Authors:** Lisa Nobis, Sanjay G. Manohar, Stephen M. Smith, Fidel Alfaro-Almagro, Mark Jenkinson, Clare E. Mackay, Masud Husain

## Abstract

Measurement of hippocampal volume has proven useful to diagnose and track progression in several brain disorders, most notably in Alzheimer’s disease (AD). For example, an objective evaluation of a patient’s hippocampal volume status may provide important information that can assist diagnosis or risk stratification of AD. However, clinicians and researchers require access to age-related normative percentiles to reliably categorise a patient’s hippocampal volume as being pathologically small. Here we analysed effects of age, sex, and hemisphere on the hippocampus and neighbouring temporal lobe volumes, in 19,793 generally healthy participants in the UK Biobank. A key finding of the current study is a significant acceleration in the rate of hippocampal volume loss in middle age, more pronounced in females than in males. In this report, we provide normative values for hippocampal and total grey matter volume as a function of age for reference in clinical and research settings. These normative values may be used in combination with our online, automated percentile estimation tool to provide a rapid, objective evaluation of an individual’s hippocampal volume status. The data provide a large-scale normative database to facilitate easy age-adjusted determination of where an individual hippocampal and temporal lobe volume lies within the normal distribution.

---

One of the most common brain imaging markers used in clinical practice to assist in the diagnosis of Alzheimer’s Disease (AD) is hippocampal volume on a structural MRI scan (Frisoni et al. 2010; Ahmed et al. 2014). Both longitudinal and cross-sectional studies have reported reduced volume of the hippocampus in patients with AD or mild cognitive impairment (MCI) compared to healthy controls (Henneman et al. 2009; Shi et al. 2009; Frankó et al. 2013). In addition, a meta-analysis of nine studies found a 3.33% difference in atrophy rate between AD and controls (Barnes et al. 2009). Increased atrophy of the hippocampus has also been associated with neurofibrillary tangle and amyloid plaque deposition, which are considered to be the hallmark features of Alzheimer’s disease (Kril et al. 2002; Schuff et al. 2009). Similarly, using the rate of hippocampal atrophy, researchers have been able to distinguish between those with MCI who progressed to AD, and those who did not (Frankó et al. 2013). Importantly, the regions of the hippocampus with the highest atrophy rate also presented with the most severe amyloid deposition (Frankó et al. 2013). Hence, hippocampal volume is considered to be a useful – and widely available – proxy to measure disease burden and progression in AD, including in clinical trials (Mielke et al. 2012; Frankó et al. 2013; Kishi et al. 2015; Choe et al. 2016).

Precise estimations of hippocampal volume may also facilitate earlier diagnosis, with some researchers reporting that the rate of hippocampal atrophy deviated from a healthy trajectory about 5.5 years before the stage of clinical AD was reached (Chételat et al. 2005). Indeed, it appears that baseline hippocampal volume and atrophy rate may be more suitable than whole brain volume to distinguish between MCI and controls (Henneman et al. 2009). It is now appreciated that estimation of hippocampal volume may also be useful for measuring disease burden or progression in other disorders. For example, hippocampal atrophy has been related to impaired episodic memory in patients with multiple sclerosis (Koenig et al. 2014) and temporal lobe epilepsy (Reyes et al. 2018), and can predict cognitive impairment in patients with Parkinson’s disease (Kandiah et al. 2014). Similarly, a cross-sectional study in patients with autoimmune encephalitis (Chakos et al. 2005) reported that the severity of the disease course is associated with lower hippocampal volume, which also predicted worse cognitive outcome (Finke et al. 2017).

In psychiatric disorders too, hippocampal volume has been implicated as an important imaging correlate. For example, reduced hippocampal volume has been repeatedly linked to depression (Campbell et al. 2004; Videbech and Ravnkilde 2004), and a meta-analysis has reported that smaller hippocampi in depressed patients predict lower response rates to antidepressant drugs (Colle et al. 2016). Similarly, in schizophrenia, hippocampal volume is reduced in chronic cases compared to healthy controls (Adriano et al. 2012). In addition, in a recent study on the effects of cognitive remediation therapy for cognitive impairment in schizophrenia, improvement was correlated with increased hippocampal volume post-treatment (Morimoto et al. 2018). These findings demonstrate that hippocampal volume estimations have the potential to provide important information to assist diagnosis, risk stratification and possibly even monitor the effects of intervention. However, several key challenges in the processing, analysis, and interpretation of structural MRI scans outside of a research setting have so far prevented this method from being used in standard clinical practice.

First, in order to calculate hippocampal volume from a structural MRI scan, the image has to be processed with software requiring expert knowledge. Automated brain segmentation tools (e.g. the FIRST tool in FMRIB Software Library (FSL) (Jenkinson and Smith 2001; Jenkinson et al. 2002), or FreeSurfer (http://surfer.nmr.mgh.harvard.edu/), and openly available standardised processing pipelines (Alfaro-Almagro et al. 2018), now make the application of this method in a clinical setting feasible.

Second, hippocampal atrophy has not only been described in pathological, but also in healthy ageing. A meta-analysis estimated the average yearly rate of hippocampal atrophy at 1.4% in healthy ageing, and 4.7% in AD (Barnes et al. 2009). Thus, in order to reliably categorise a patient’s hippocampal volume, clinicians and researchers require access to age-related normative percentiles to distinguish between healthy and pathological ageing. Few studies have attempted to provide such normative values – or nomograms – not least because of methodological obstacles. The acquisition of large MRI datasets is costly, time consuming, and involves highly specialised hardware and staff. Previously published studies on hippocampal volume in the general population are therefore often limited by small sample sizes, low statistical power, and cohort effects (Ioannidis 2011; Button et al. 2013; Fraser et al. 2015; Nord et al. 2017). In addition, other potentially confounding factors, such as method used, sex, head size, years of education, smoking status, and effects of normal ageing have to be considered when applying such normative percentiles.

Here we report a first attempt to provide detailed normative information on the hippocampus and neighbouring temporal lobe volumes in relation to age, sex, total grey matter volume, and symmetry in 19,793 generally healthy older adults in the UK Biobank resource (http://www.ukbiobank.ac.uk). We chose a model-free sliding window approach to study the relationships of temporal lobe volumes and total grey matter with age, as this method makes few assumptions and does not impose a linear relationship. While efforts to harmonize analysis across MRI datasets have not yet been successful, the large dataset analysed here allows an attempt to provide robust normative data when these are used with estimates obtained using the same, or similar, acquisition protocols, hardware and pre-processing pipelines.

We also supply a webtool, where a patient’s estimated brain volume may be entered to calculate their hippocampal volume status as compared to age-matched norm values (http://www.smanohar.com/biobank). These norms will become especially useful with the acquisition of longitudinal health outcomes of participants in UK Biobank. The UK Biobank cohort includes over 500,000 participants aged 40-69 years, of which a subset of 100,000 participants will undergo brain imaging. Within this subset, 6000 participants are likely to have developed AD by 2027 (Miller et al. 2016). UK Biobank has been granted access to the UK National Health Service records and can thus follow up on future health outcomes of participants, making it a powerful resource to study disease progression. In addition, both the acquisition and analysis of MRI are standardised across the scanning sites according to publicly available protocols designed by the UK Biobank Imaging Working Group (www.ukbiobank.ac.uk/expert-working-groups). This increases generalisability of results and resolves some of the issues of previous studies using different scanners and protocols.

## Methods

### Participants

The most recent release of brain imaging data of the UK Biobank includes scans of 20,542 participants aged 45-80 years at the time of scanning. Participants with self-reported neurological and psychiatric disorders, substance abuse disorders or a history of head trauma were excluded from our analyses, leaving a sample of 19,793 generally healthy participants scanned in 2014-2018.

### MRI acquisition and analysis

Image acquisition and pre-processing of MRI scans have been performed by UK Biobank. Brain images were acquired on a Siemens Skyra 3.0 T scanner (Siemens Medical Solutions, Germany) with a 32-channel head coil. T1-weighted images with 1 mm^3^ isotropic resolution were previously analysed with FMRIB Software Library (FSL) (http://fsl.fmrib.ox.ac.uk/fsl), with image-derived phenotypes (IDPs – imaging summary statistics such as brain volume and hippocampal volume) made available for general access. More detailed information on MRI acquisition and analysis have been reported elsewhere (Miller et al. 2016; Alfaro-Almagro et al. 2018). UK Biobank also published a standardised MRI analysis pipeline (FMRIB’s Biobank Pipeline version 1.0) that is freely available to the public, including the source code (https://git.fmrib.ox.ac.uk/falmagro/UK_biobank_pipeline_v_1) (Alfaro-Almagro et al. 2018).

### Statistics

All calculations were performed in MATLAB 2017b with the exception of Joinpoint regression, which was computed with the Joinpoint Regression Program (version 4.6.0.0; Surveillance Research Program, National Cancer Institute) (Kim et al. 2000).

#### Data preparation

Outliers in brain volume estimations were identified and excluded based on a Median Absolute Deviation of > 5. Brain volumes were always corrected for scanning date, and when indicated, for head size and age. This was done with a General Linear Model, ‘regressing out’ variance accounted for by the confounds. Head size was corrected for using a head scaling variable obtained from the MRI scan, which is the volumetric scaling for the transformation of the native head image to standard space, with final scaling being driven by outer-skull surface (Smith et al. 2002, 2004). A number of studies have shown that global and regional brain volumes correlate positively with head size (Barnes et al. 2010). Thus, when investigating sex differences in brain volume, head size should be corrected for. This was done by regression rather than using a ratio measure (e.g. brain volume/head size), as not all brain volumes scale proportionately with head size (Barnes et al. 2010). However, measures of head size related variables vary widely in the literature (e.g. total intracranial volume, total brain volume, MRI head scaling, or body height), leading to heterogenous results. We therefore report both head size corrected and un-corrected results.

#### Analysis

For comparison we also present analyses for volumes of some structures near the hippocampus within the temporal lobe. Available cortical grey matter volume IDPs within the UK Biobank dataset that are reported here include superior temporal gyrus, middle temporal gyrus, inferior temporal gyrus, fusiform gyrus, parahippocampal gyrus and temporal pole. T-tests (with Bonferroni-correction for resulting p-values) were calculated for sex differences in age and brain volume (unpaired t-test), as well as for differences in hemisphere (paired t-test), hypertension status (unpaired t-test), and hippocampal volume estimations by method (FSL-FIRST, vs. FSL-FAST within Atlas-based regions of interest; paired t-test). While FSL-FIRST is a segmentation/registration tool that estimates subcortical volumes based on shape and appearance models (Patenaude et al. 2011), FSL-FAST is a brain tissue segmentation tool (Zhang et al. 2001), with which subcortical volumes were estimated from grey matter within Atlas-based regions of interest. ANOVAs were run for the effects of level of education and smoking status on brain volumes. Education level was binarized into higher education degree (college/university degree, NVQ), and lower education degree (A levels, O levels, CSE). For visualisation purposes, volume differences based on sex, hemisphere, hypertension status, and education are plotted as proportional differences, calculated with

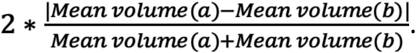

All MATLAB analysis code will be made available on reviewer request or publication.

#### Normative Percentiles

The 2.5^th^, 5^th^, 10^th^, 25^th^, 50^th^, 75^th^, 90^th^, 95^th^, and 97.5^th^ percentiles for hippocampal volume (left and right) and total grey matter were calculated separately for males and females. This was done using residuals from a corresponding sliding-window analysis (see below), that were added back onto the mean of the brain volume estimations. These values were then used as the age-adjusted input to calculate the percentiles.

#### Sliding-window curves for relationship of volume and age

To analyse trajectories and quantiles of measured volumes as a function of age, we applied a sliding-window, model-free analysis, in which an age-window of observations of *fixed age-quantile width* was moved along the age distribution (based on function ‘conditionalPlot’ in https://osf.io/vmabg/; (Manohar 2019). The windows overlapped, so that each window contained 10% of the participants. The analysis of interest was performed for each window, providing in this case the mean volume of brain structure, such as hippocampal volume. The volumes were then smoothed using the moving average method with a gaussian kernel of 20. Results from using varying smoothing kernels and quantile widths can be found in the **Supplementary Material**. The mean brain volume was plotted against the mean bin centre of age, along with standard errors of the mean. This provided an estimate of mean brain volume that varied smoothly as a function of age. This method enables representative contours to be calculated for each quantile as a function of age, in a non-parametric, data-driven manner. Here, we present sliding-window curves across age for all available temporal lobe areas, and total grey matter volume. In addition, curves for the ratio of the individual temporal lobe volumes to the rest of grey matter volume are shown, as well as the slope of the hippocampal volume versus age (in **Supplementary Material**), using bootstrapped confidence intervals.

We tested for differences in the peak of the ratio curves between males and females. This was performed using a permuted t-statistic, correcting p-values for false discovery rate. Specifically, we permuted matched age bins of the male and female groups, i.e. shuffling males and females within age-windows. We then calculated the p-value as the proportion of permuted datasets (n = 5000), which produced a mean difference between the peaks of the curves at least as extreme as the one observed from the actual data.

This data-driven analysis is particularly useful for large samples. As opposed to traditional curve fitting methods, it makes few assumptions, has only one free parameter (number of samples per bin), and improves visualisation and interpretation of results and corresponding errors. Note that as there are fewer participants at the minimum and maximum age limits compared to around the mean age, this method leads to wider age ranges for the lowest and highest percentile bins. As shown in **Supplementary Figure S12**, similar effects to our main analysis were observed when using windows of fixed age bins (5 years width), rather than bins of fixed numbers of observations. Again, the mean brain volume was computed within each age bin, and plotted against the mean bin centre of that bin, along with standard errors.

#### Joinpoint regression

To determine if there were significant points at which the rate of hippocampal atrophy changed across age, we used joinpoint regression. This is a regression analysis recommended by the National Cancer Institute (NCI, https://surveillance.cancer.gov/joinpoint/, (Kim et al. 2000)) to estimate time points of change in trend data, often used for changes in cancer incidence rates. The analysis fits separate line segments to trend data, which are connected at positions called joinpoints. No joinpoints would suggest no changes in slope, and thus no changes in, for example, cancer incidence. Here we applied joinpoint regression to test for the presence of one or more joinpoints (i.e. inflection points) along the brain volume trajectories across age. Thus, instead of cancer incidence rate, here the dependent variable was mean volume per 1-year, non-overlapping age window. Bins of 1-year width were chosen to retain adequate accuracy with a feasible number of datapoints. Similarly, instead of diagnosis year, the independent variable used here was age group.

The Joinpoint Regression Program fits between zero and a chosen maximum number of joinpoints to the data. Starting from zero joinpoints, we tested whether more joinpoints are statistically significant (*p* < .05) using a Monte Carlo Permutation method, assuming uncorrelated errors. The maximum was chosen at 4 joinpoints, as recommended for our number of data points (Kim et al. 2000). This was done with hippocampal volume, rest of total grey matter volume (total grey matter minus hippocampal volume), superior temporal gyrus, middle temporal gyrus, inferior temporal gyrus, fusiform gyrus, parahippocampal gyrus, and temporal pole volume, for males and females separately. More detailed information on methods used within the Joinpoint Regression Program can be found at https://surveillance.cancer.gov/help/joinpoint/.

## Results

### Demographics

Demographics as well as means and standard deviations for uncorrected hippocampal volume, with respect to head size, by hemisphere and sex are summarised in Table 1.

**Table 1:**
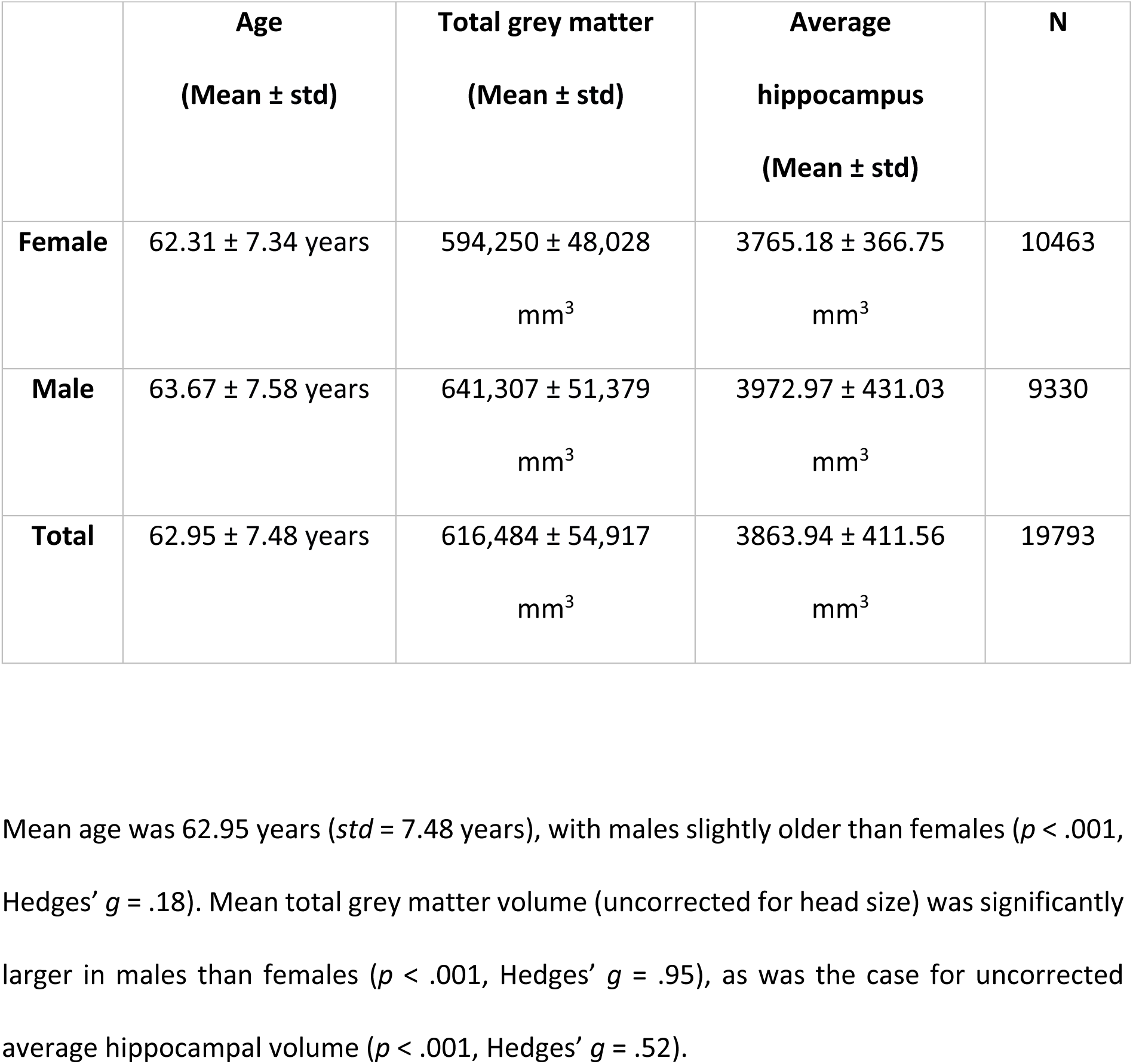
Demographics of sample

### Effects of hypertension, smoking, higher BMI and lower education on brain volume

Before further analysing the demographic information, brain volumes were corrected for head size as described above. As effects of hypertension status, Body-Mass-Index (BMI), education, and smoking status on brain volumes may also be confounded by age, variance introduced by age was also regressed out for the following results on effects of hypertension, BMI and education only.

Participants with self-reported hypertension had significantly lower hippocampal (*p* < .001, Hedges’ *g* = .06) and total grey matter volumes (*p* < .001, Hedges’ *g* = .17), than participants without self-reported hypertension, though with small effect sizes (**Suppl. Figure S1**). There was a significant, but very small negative correlation between BMI and hippocampal volume (*r* =-.03, *p* < .001, *n* = 18,653), as well as between BMI and total grey matter volume (*r* =-.11, *p* <. 001, *n* = 18,653), with larger BMI being weakly associated with smaller volumes. There was also a small significant positive effect of level of education on average hippocampal volume (*p* < .001, Hedges’ *g* = .07, *n* = 19,646). Higher education levels were associated with larger volumes than lower level education levels. There was no significant effect of education on total grey matter volume (*p* = .77, *n* = 19,646, **Suppl. Figure S2**).

Hippocampal (*F* = 8.26, *p* < .001, *n* = 19,646) and total grey matter volume (*F* = 60.12, *p* < .001, *n* = 19,646) were significantly smaller for current smokers than for people who have never smoked, and people who have previously smoked. In addition, previous smokers had larger hippocampal and total grey matter volumes than current smokers (**Suppl. Figure S3**).

**Supplementary Tables 1-3** present data for means, standard deviations, and significance tests of hypertension, BMI, education level, and smoking status for all temporal lobe volumes as well as total grey matter volume. Percent differences for all temporal lobe volumes are shown in **Supplementary Figures S1-3**.

### Normative percentiles

Example nomograms for the 2.5^th^, 5^th^, 10^th^, 25^th^, 50^th^, 75^th^, 90^th^, 95^th^, and 97.5^th^ percentiles for left and right hippocampal volume in women and men, uncorrected for head size, are shown in Figures 1-4. Overlapping, age-adjusted windows of 10% of observations are plotted, with age on the x-axis corresponding to the median age of participants within the overlapping quantile windows. Because there are fewer observations within particularly young (starting at 45 years) and particularly old (up to 80 years) participants, quantile bins for the youngest and oldest participants are wider, and thus have a median age around 50 and 75 years, respectively. Additional moving average smoothing with a gaussian kernel of 20 resulted in percentiles plotted across ages 51 years to 72 years (52 years to 73 years for males). The nomograms are provided separately for females and males as the previous analyses have shown differential sex by age trajectories. Additional nomograms split by hemisphere and sex for total grey matter, as well as plots with volumes corrected for head scaling can be found in the Supplementary Material (**Suppl. Figures S4-11**).

**Figure 1:**
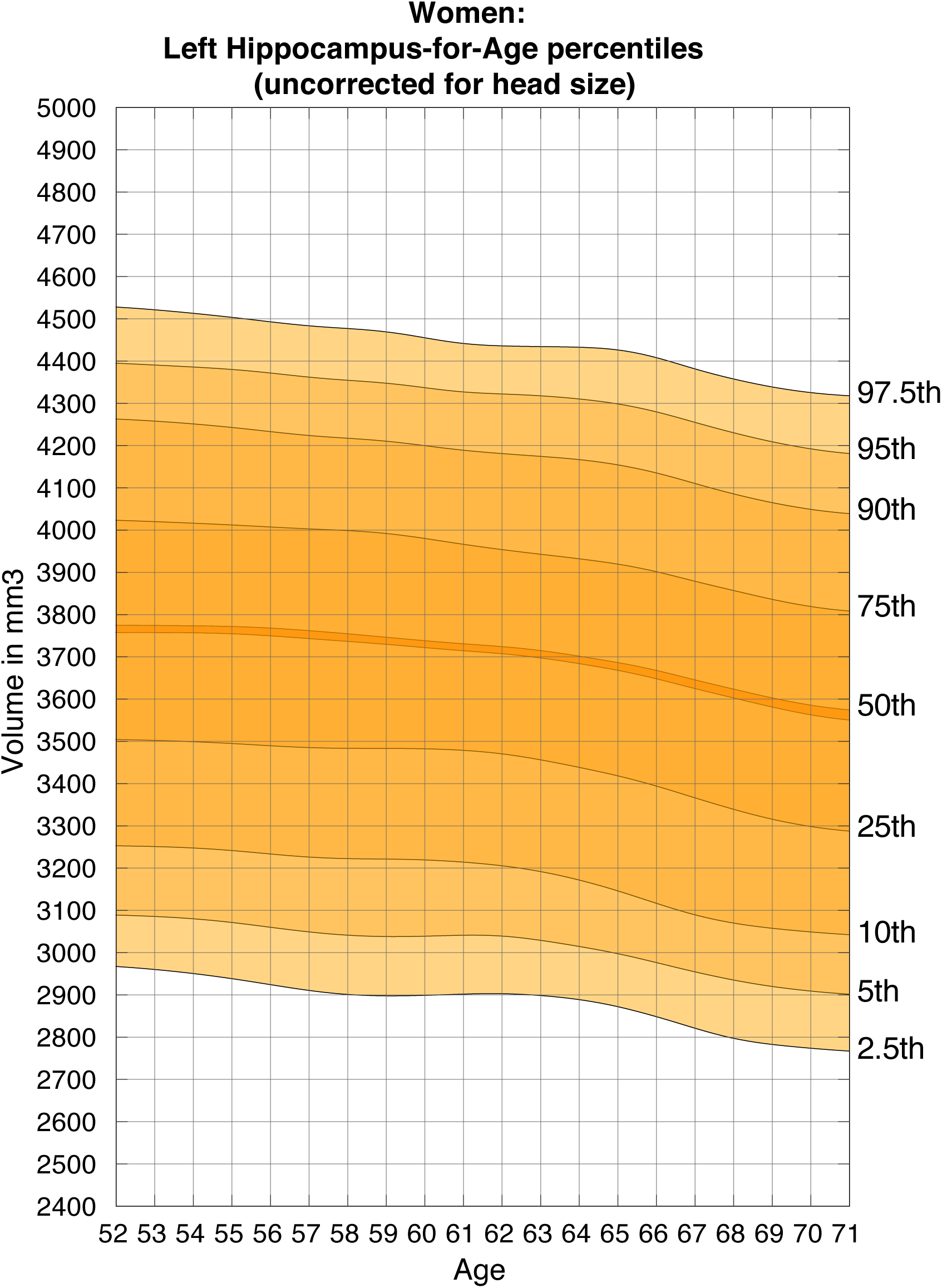
Nomogram of uncorrected left hippocampal volume for females. Figures show the quantiles of hippocampal volume for the group of individuals in each age window. The x-axis indicates the median age of the window. The mid line indicates the median. For someone with a given age and volume, a percentile can be read off the chart, indicating the proportion of the Biobank cohort who have a hippocampal volume below that of the person.

**Figure 2:**
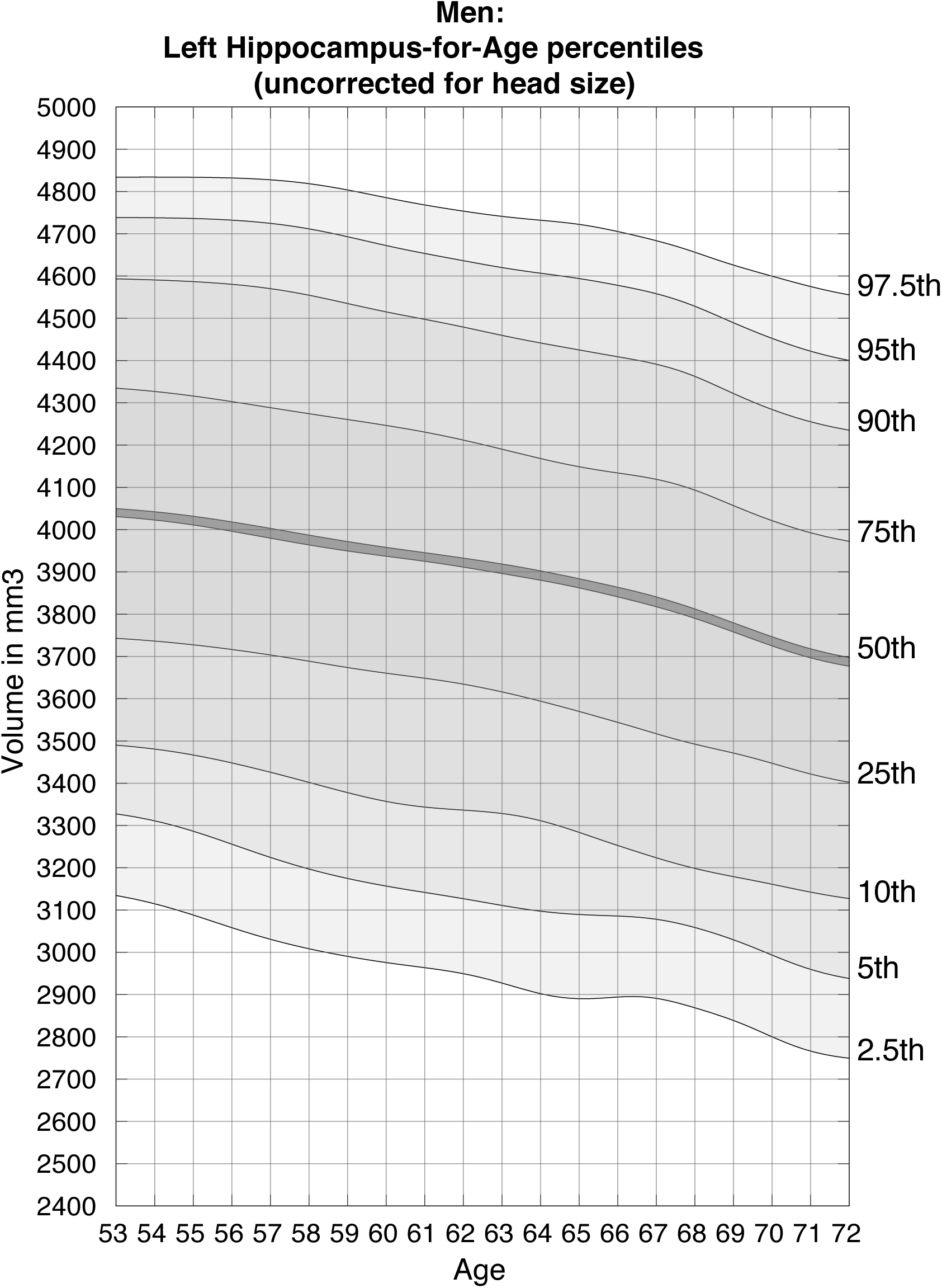
Nomogram of uncorrected left hippocampal volume for males. Figures show the quantiles of hippocampal volume for the group of individuals in each age window. The x-axis indicates the median age of the window. The mid line indicates the median. For someone with a given age and volume, a percentile can be read off the chart, indicating the proportion of the Biobank cohort who have a hippocampal volume below that of the person.

**Figure 3:**
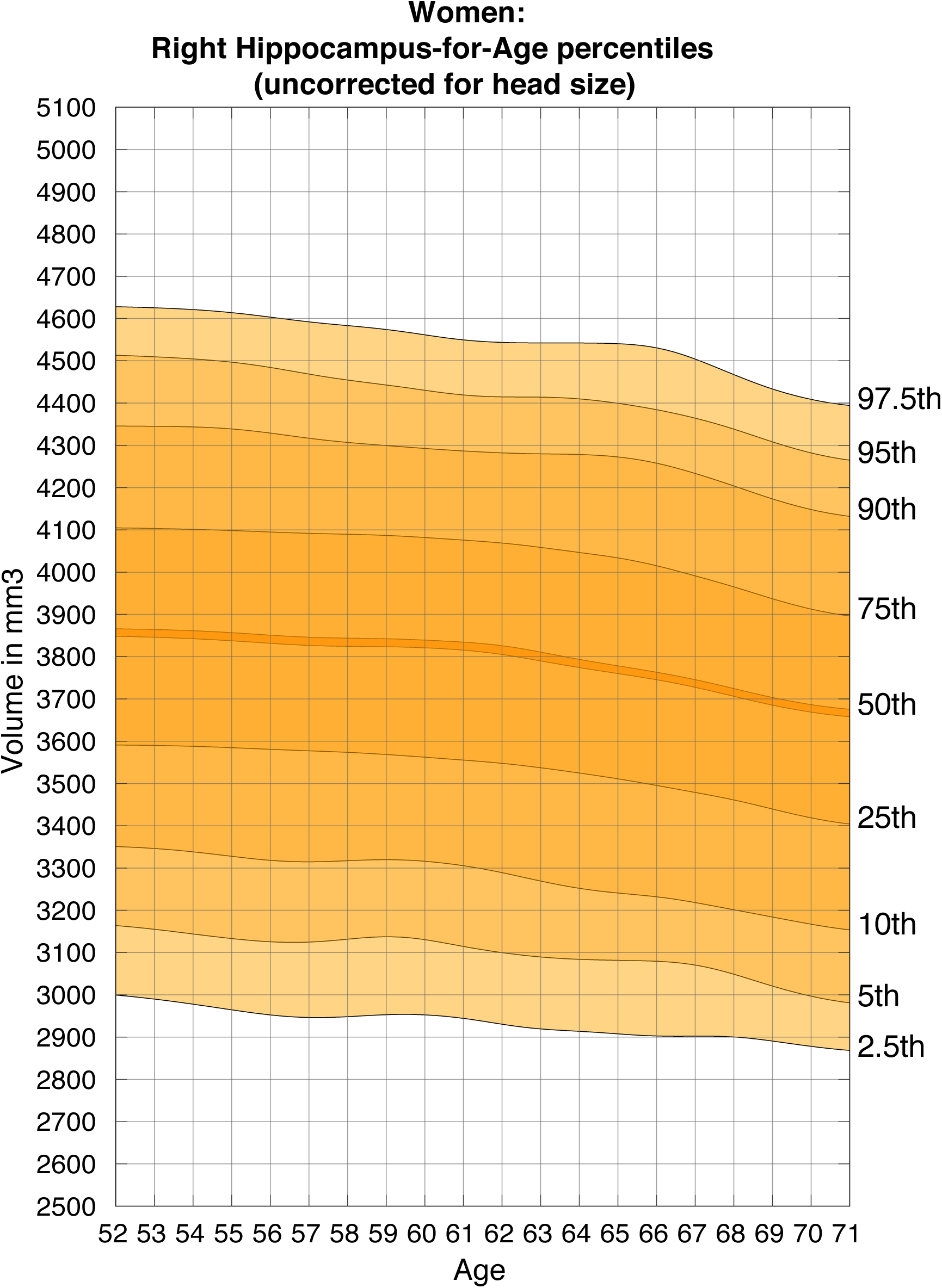
Nomogram of uncorrected right hippocampal volume for females. Figures show the quantiles of hippocampal volume for the group of individuals in each age window. The x-axis indicates the median age of the window. The mid line indicates the median. For someone with a given age and volume, a percentile can be read off the chart, indicating the proportion of the Biobank cohort who have a hippocampal volume below that of the person.

**Figure 4:**
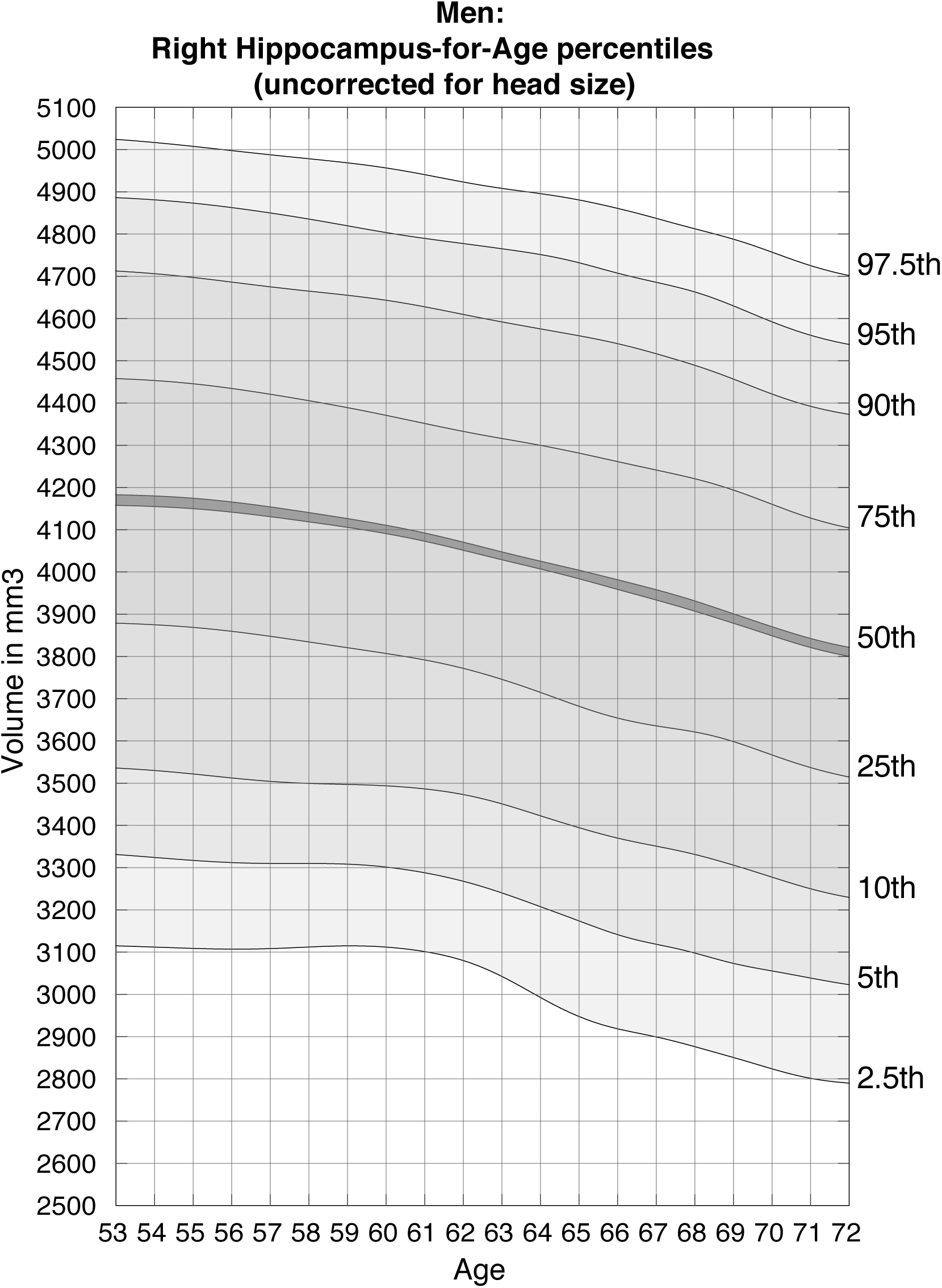
Nomogram of uncorrected right hippocampal volume for males. Figures show the quantiles of hippocampal volume for the group of individuals in each age window. The x-axis indicates the median age of the window. The mid line indicates the median. For someone with a given age and volume, a percentile can be read off the chart, indicating the proportion of the Biobank cohort who have a hippocampal volume below that of the person.

The provided nomogram webtool (http://www.smanohar.com/biobank/) enables clinicians to quickly calculate a patient’s percentile by entering the volume estimation from the automated analysis pipeline. For example, a female aged 64 years with a left hippocampal volume of 3.15 cm^3^ (corrected for head size) would score within the 5^th^ percentile. This means that 95% of the women closest to her age in the current sample had a larger volume, highlighting potentially pathological atrophy and providing additional information for the diagnosing clinician.

### Brain volume characteristics

#### Temporal lobe volumes differed between hemispheres

A paired t-test resulted in a significant difference between right and left hippocampal volume (uncorrected for head size) with *right larger than left* volume asymmetry (**see Figure 5** for percent difference, *p* < .001, *Hedges’ G* = .24). The same effect was found in the superior temporal gyrus (*p* < .001, *Hedges’ G* = .80) and inferior temporal gyrus (*p* < .001, *Hedges’ G* = .12). On the other hand, there was a significant *left larger than right* volume asymmetry for the fusiform gyrus (*p* < .001, *Hedges’ G* = 1.30) and parahippocampal gyrus (*p* < .001, *Hedges’ G* = .48). Asymmetry effects of the same direction in the temporal pole were also significant, however with a very small effect size (*Hedges’ G* = .04). There was no significant difference between left and right hemisphere for middle temporal gyrus (p = .61). See **Supplementary Table 4** for means, standard deviations, and significance tests for all temporal lobe volumes. Head size was not corrected for as left/right differences are within-subject, and will be largely unaffected by head size. There was no correlation between handedness and asymmetry for any of the tested volumes (calculated by asymmetry = left volume – right volume; all *p* > α/7 = .007).

**Figure 5:**
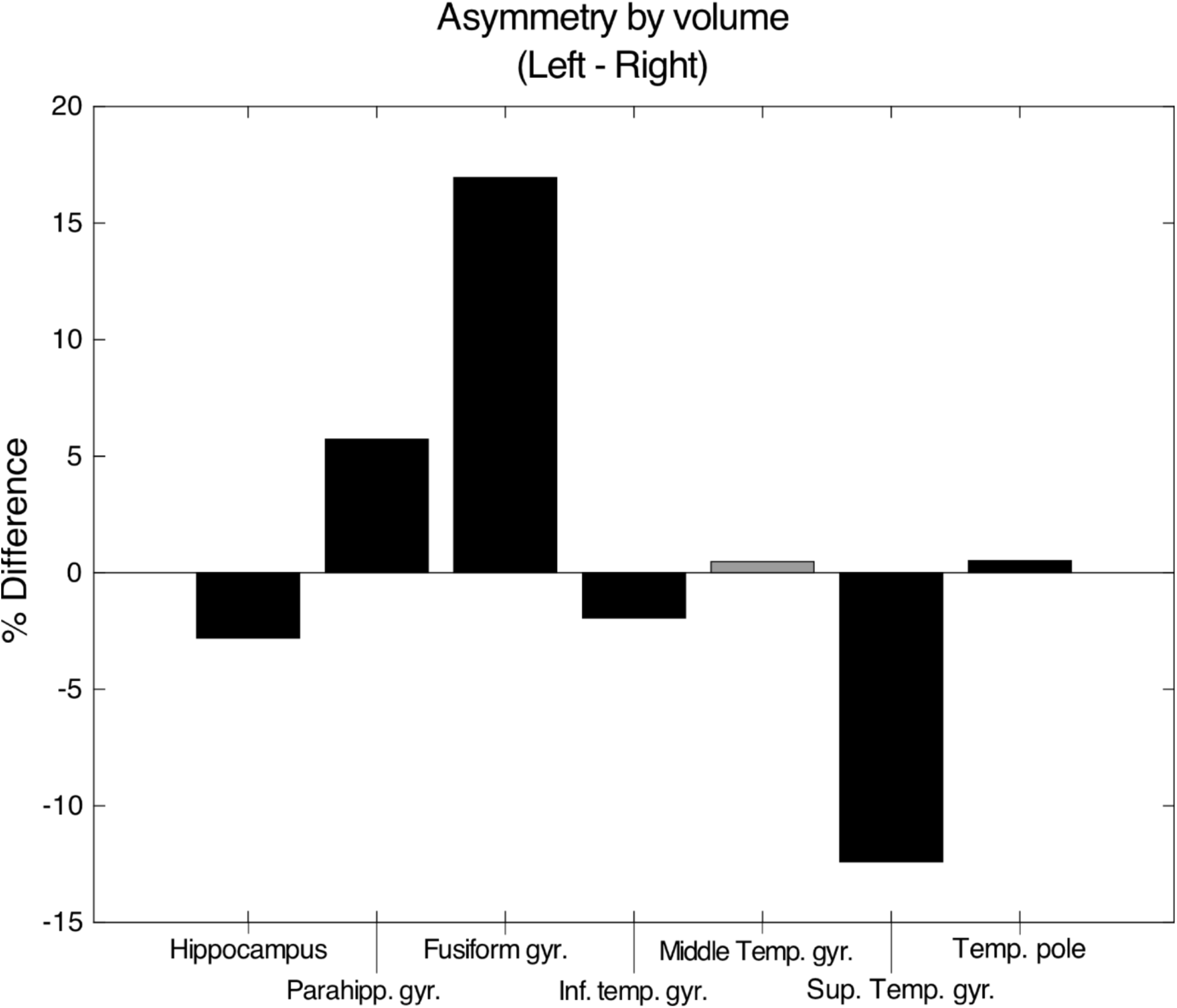
Percent differences between left and right volumes. Negative percentage differences correspond to larger right volumes, positive percent differences correspond to larger left volumes. Percent differences are shown for visualisation, black bars indicate a significant corresponding t-test for absolute volume differences at *p < .001 (Bonferroni-corrected α = .05/8 = .006)*.

#### Temporal lobe volumes by sex

Means, standard deviations, and significance tests for all brain volumes by sex, corrected for head size and age, can be found in **Supplementary Table 5.** There was no significant difference of mean (average) hippocampal volume between males and females (*p* > .05). Sex differences for the remaining temporal lobe volumes, and for total grey matter volume were significant in different directions, though effect sizes were mostly very small (See **Figure 6** for percent differences). Small-to-medium effect sizes were seen in total grey matter volume (females larger than males), parahippocampal gyrus (males larger than females), and temporal pole (males larger than females).

**Figure 6:**
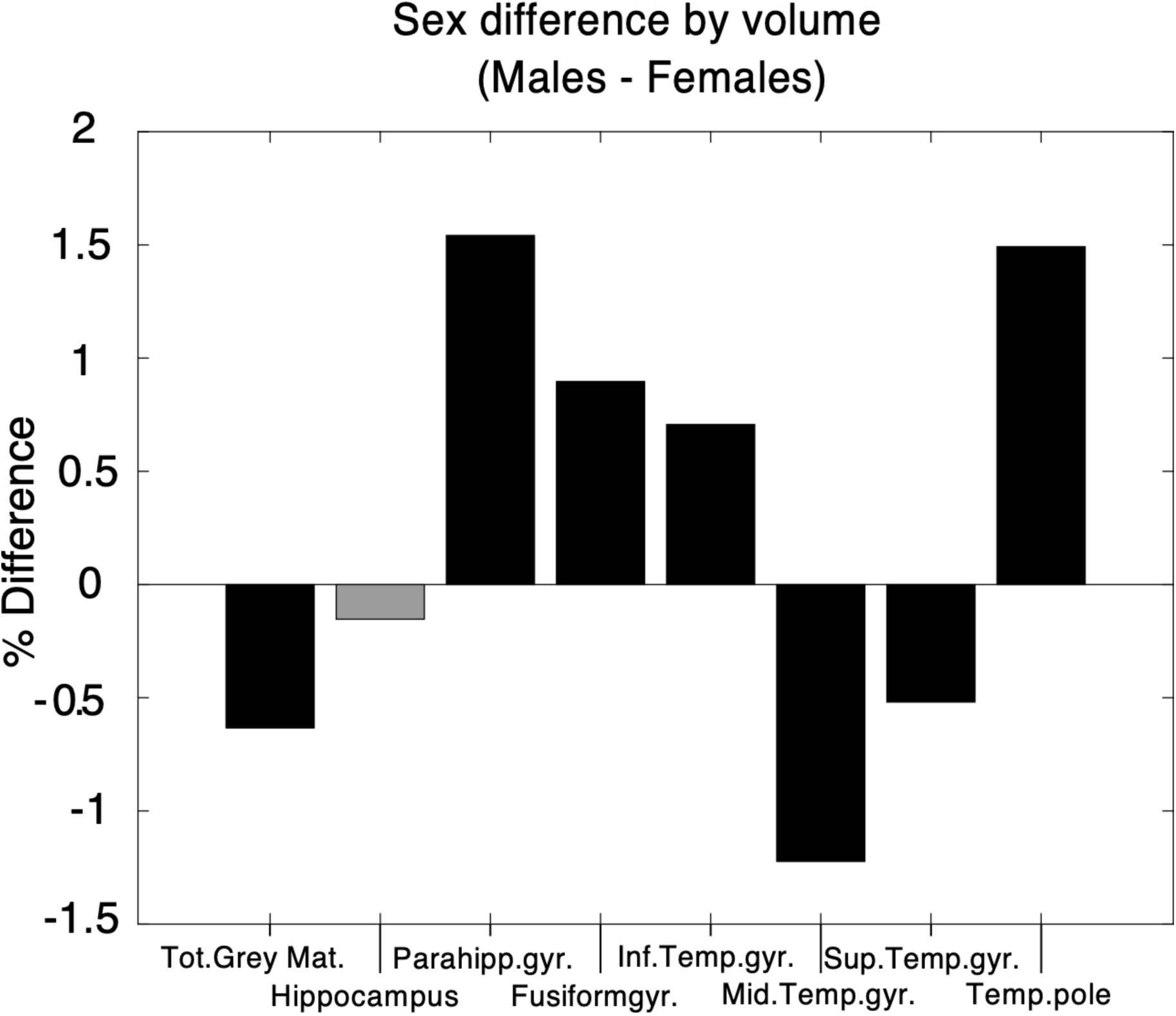
Percent differences between males and females. Positive percent differences correspond to larger volumes in males, negative percent differences correspond to larger volumes in females. Black bars indicate a significant corresponding t-test for absolute volume differences at *p < .001 (Bonferroni-corrected α* = *.05/8 = .006)*.

#### Method of estimation did not alter Hippocampal volume

Mean hippocampal volumes differed between methods. On average FSL-FAST estimates (*mean* = 4285.08 mm^3^, *std* = 309.32 mm^3^) were larger than estimates calculated with FSL-FIRST (*mean* = 3863.40 mm^3^, *std* = 353.61mm^3^, *p* < .001, Hedges’ *g* = 1.27). Estimates were corrected for head size, scanning date, and age. However, there were no differences in the overall results from the joinpoint and sliding-window analyses between the two hippocampal volume estimates (see sections below). As FSL-FIRST is the recommended and most widely used method for subcortical segmentation, we present analyses throughout using the IDPs that were generated using FSL-FIRST.

#### Trajectory of mean hippocampal volume with age demonstrate an inflection

We plot mean hippocampal volume, total grey matter volume (both corrected for head size), and hippocampal volume as a percentage of grey matter volume as a function of age. To plot this we used a sliding-window method, using windows of 10% of the population (**Figure 7**). Age on the x-axis corresponds to the median age of participants within the overlapping quantile windows. Because there are fewer observations within particularly young and particularly old participants, the median ages of the quantile bins for the youngest and oldest participants are around 50 and 75 years, respectively. Plots for left and right hippocampal volumes, as well as plots for fixed 5-year age-bins rather than fixed quantiles for bilateral hippocampal volume, are provided in the Supplementary Material (**Suppl. Figures S12-13**).

**Figure 7:**
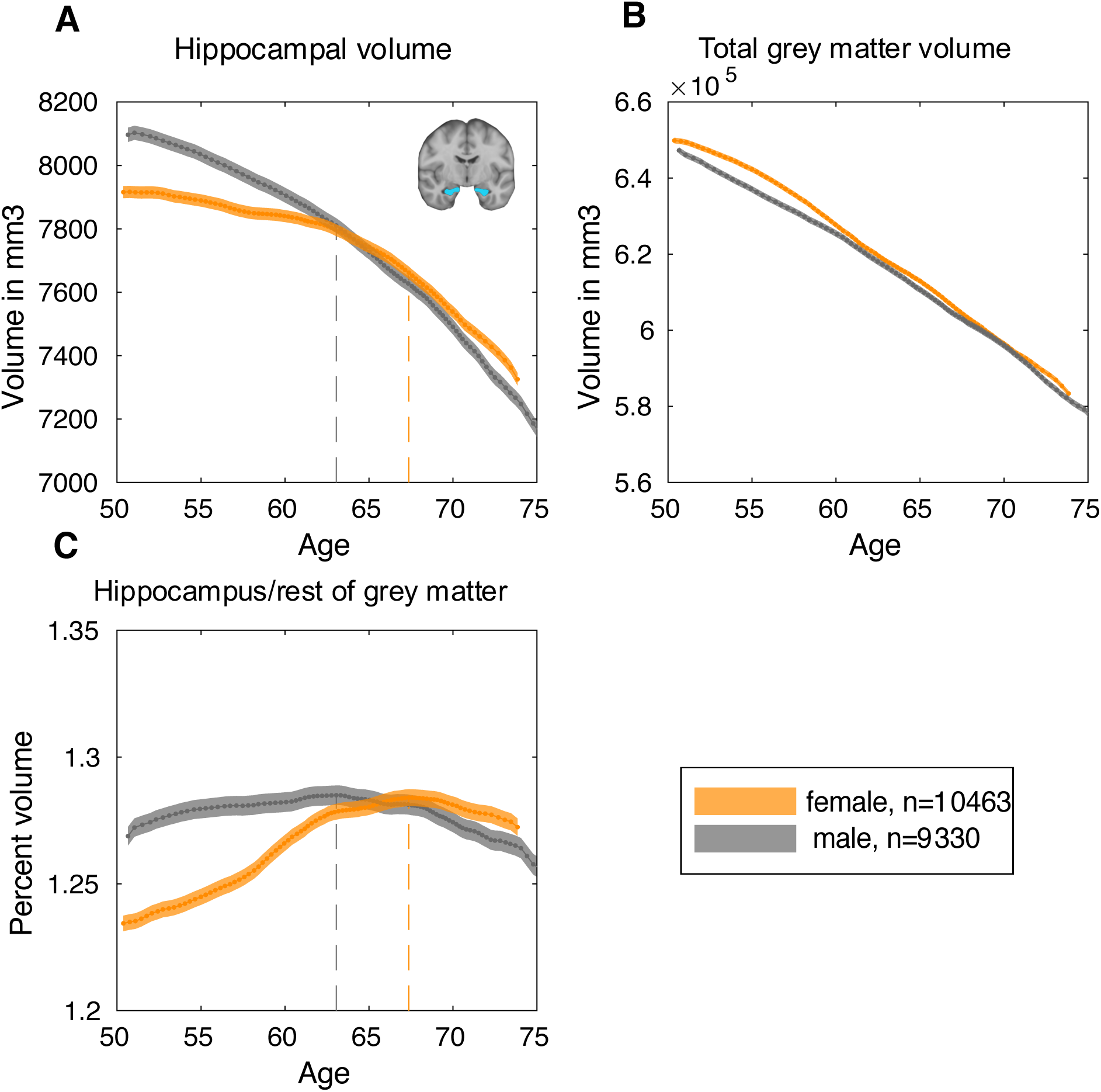
Trajectory of hippocampal volume with age. Dashed lines indicate points of maximum ratio. **A**. Mean bilateral hippocampal volume including standard errors as a function of age, corrected for head size. **B**. Mean total grey matter volume including standard errors as a function of age, corrected for head size. **C**. Mean hippocampal volume to rest of grey matter ratio including standard errors as a function of age.

Whereas whole brain volume appears to decline linearly as a function of age for both men and women (**Figure 7B**), in women the rate of hippocampal volume loss increases at ~60 years (**Figure 7A**). This inflection point (i.e. maximum slope), can be calculated by numerical differentiation, as shown in **Supplementary Figure S14** (females: bin(*max slope)* = 58.67 – 61.28 years, max slope = −0.27; males: bin(*max slope)* = 44.56 – 52.73 years, max slope = −2.52).

To compare hippocampal volume more directly to the rest of the grey matter, the ratio between these two volumes was calculated. In females, the ratio of hippocampal volume to the rest of grey matter (**Figure 7C**) peaks around age 67 years (bin(*max ratio)* = 66.33 years – 68.44 years), suggesting that the rate of volume loss in the hippocampus is smaller relative to the rest of grey matter. As age increases further, this pattern is then reversed. In males the ratio of hippocampal volume relative to the rest of grey matter peaks around age 63 years (bin(*max ratio)* = 61.69 years – 64.32 years), before declining thereafter. These sex differences in the age of transition (maximum ratio) were significant. Males reached peak hippocampal ratio at a significantly younger age than females, calculated by permuting matched age bins of the male and female groups to compute the null distribution of the difference in the peak (*n(permutations)* = 5000*, p* < .001). This p-value represents the proportion of permuted datasets that produced a mean difference between the peaks of the curves at least as extreme as the one observed from the actual data. These results were replicated using no smoothing **(Suppl. Fig S15)**, a smoothing kernel of 10 rather than 20 **(Suppl. Fig S16)**, as well as quantile bins with 25% **(Suppl. Fig S17-S18)** instead of 10% of participants (range of median age at ratio maxima for females: 67.37 – 68.21 years; for males: 62.48 – 63.32 years).

In order to confirm that there was a significant change in slope of the trajectory of hippocampal volume, joinpoint regression (Kim et al. 2000) was applied. This method tests statistically for the presence of zero, or up to four inflection points (joinpoints) in the slope of mean hippocampal volume across age. The maximum number of joinpoints tested for is selected based on the number of observations. There was one inflection point for both males and females (Figure 8A-B). In females, there was a significant change in slope at age-group 64-65 years (change of slope Δm = −32.44mm^3^; age 95% CI [61-62 years, 67-68 years], *p* <. 0001). In males, the change in slope was significant at age-group 63-64 years (change of slope Δm = −26.45m 95% CI [58-59 years, 66-67 years], *p* < .0001; see Table 2 for Model estimates).

**Table 2:** Model Estimates from joinpoint regression of hippocampal volume

|  | **Female** | **Male** |
| --- | --- | --- |
| **DF** | 21 | 21 |
| **Joinpoint** | 64-65 years | 63-64 years |
| **Joinpoint 95% LCL** | 61-62 years | 58-59 years |
| **Joinpoint 95% UCL** | 67-68 years | 66-67 years |
| **Slope Change Estimate** | -32.44 mm<sup>3</sup> | -26.45 mm<sup>3</sup> |
| **Slope Change Std Error** | 4.04 mm<sup>3</sup> | 4.94 mm<sup>3</sup> |
| **Slope Change Test Statistic\*** | -8.03 | -5.35 |
| **Slope Change p-value** | <.0001 | < .0001 |

\* the slope change test statistic is the parameter estimate divided by the standard error that follows a t distribution with 21 degrees of freedom.

**Figure 8:**
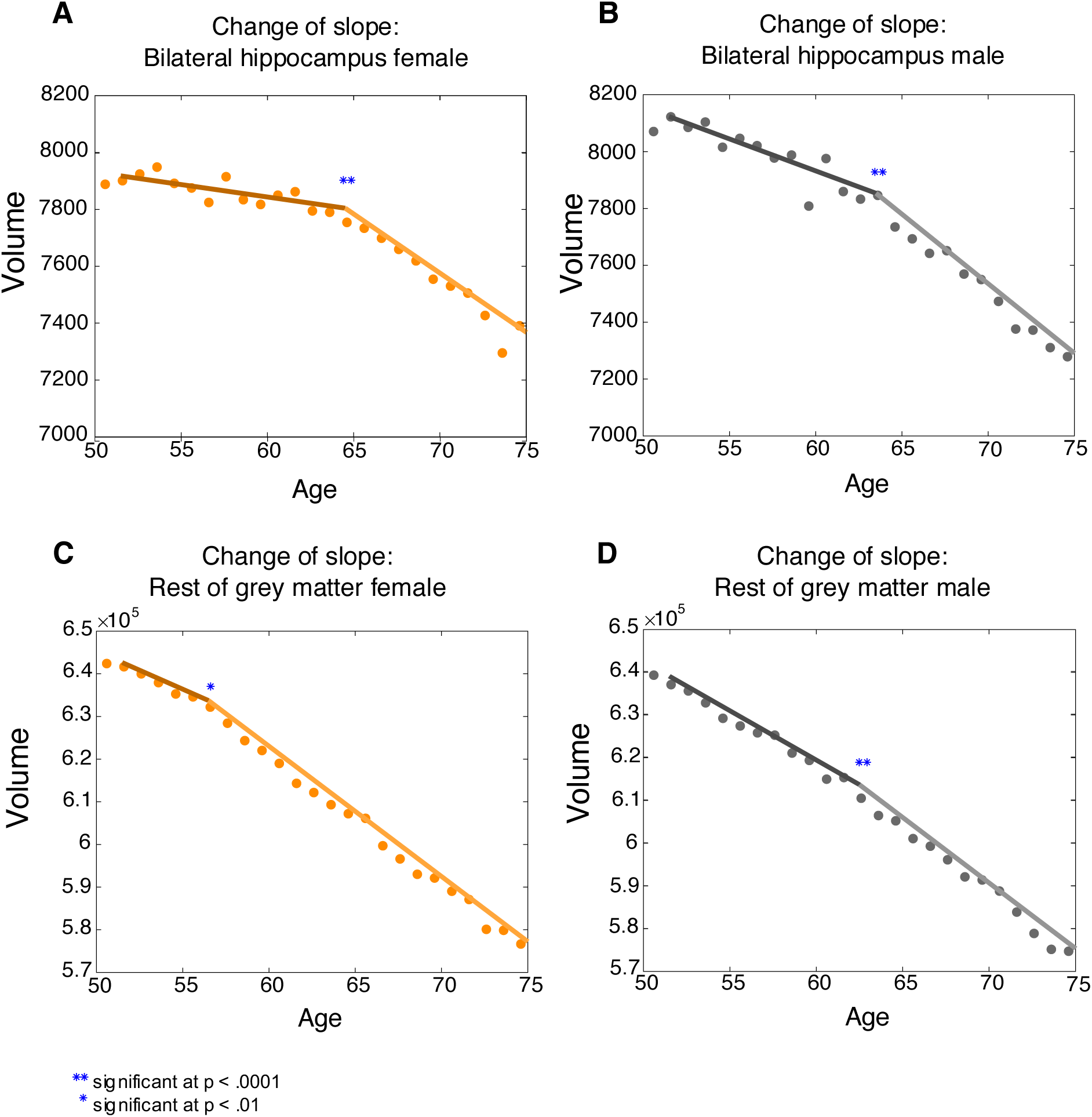
Joinpoint analysis for hippocampal and total grey matter volume. **A**. Joinpoint (change of slope) in bilateral hippocampal volume over age in females **B.** Joinpoint bilateral hippocampus in males. **C.** Joinpoint in rest of grey matter volume (total grey matter – bilateral hippocampus) in females **D**. Joinpoint in rest of grey matter for males.

For the rest of the grey matter, there was one joinpoint in the slope (total grey matter – bilateral hippocampus) across age for females (Figure 8C), with a significant change in slope of Δm = −1273.98mm^3^ at age-group 56-57 years (*t*(21) = −2.98, 95% CI [53-54 years, 58-59 years], *p* =.007). By contrast, there was also a small, but significant change in slope for the rest of grey matter across age in males at age-group 62-63 years (Δm = −754.14mm^3^, *t*(21) = −4.87, 95% CI [57-58 years, 66-67 years], *p* < .0001) (Figure 8D).

#### Trajectories of other temporal lobe volumes with age

Next, we aimed to demonstrate that these inflections were specific to the hippocampus. Mean volumes (corrected for head size) were plotted for the remaining brain areas within the temporal lobe available from UK Biobank, namely parahippocampal gyrus, fusiform gyrus, superior temporal gyrus, middle temporal gyrus, inferior temporal gyrus, and temporal pole (Figure 9). The same sliding-window and joinpoint regression methods were used. All these temporal lobe regions show a largely linear negative relationship with age. In keeping with this, and in contrast to the findings for the hippocampus, there were no significant joinpoints or changes in slope for either males or females in superior temporal gyrus, middle temporal gyrus, inferior temporal gyrus and fusiform gyrus (*p* > .05). However, there was a significant joinpoint for both males and females in parahippocampal gyrus volume, similar to the joinpoints found for the hippocampus (females: Δm = −26.97mm^3^ at age-group 66-67 years, *t*(21) = −5.40, 95% CI [62-63 years, 70-71 years], *p* < .0001; males: Δm = −30.81mm^3^ at age-group 59-60 years, *t*(21) = −3.77, 95% CI [56-57 years, 64-65 years], *p* = .001) (Figure 8D). There was also a significant joinpoint in volume of temporal pole in males, at age-group 55-56 years (Δm = −76.97mm^3^, *t*(21) = −2.57, 95% CI [53-54 years, 57-58 years], *p* =. 02).

**Figure 9:**
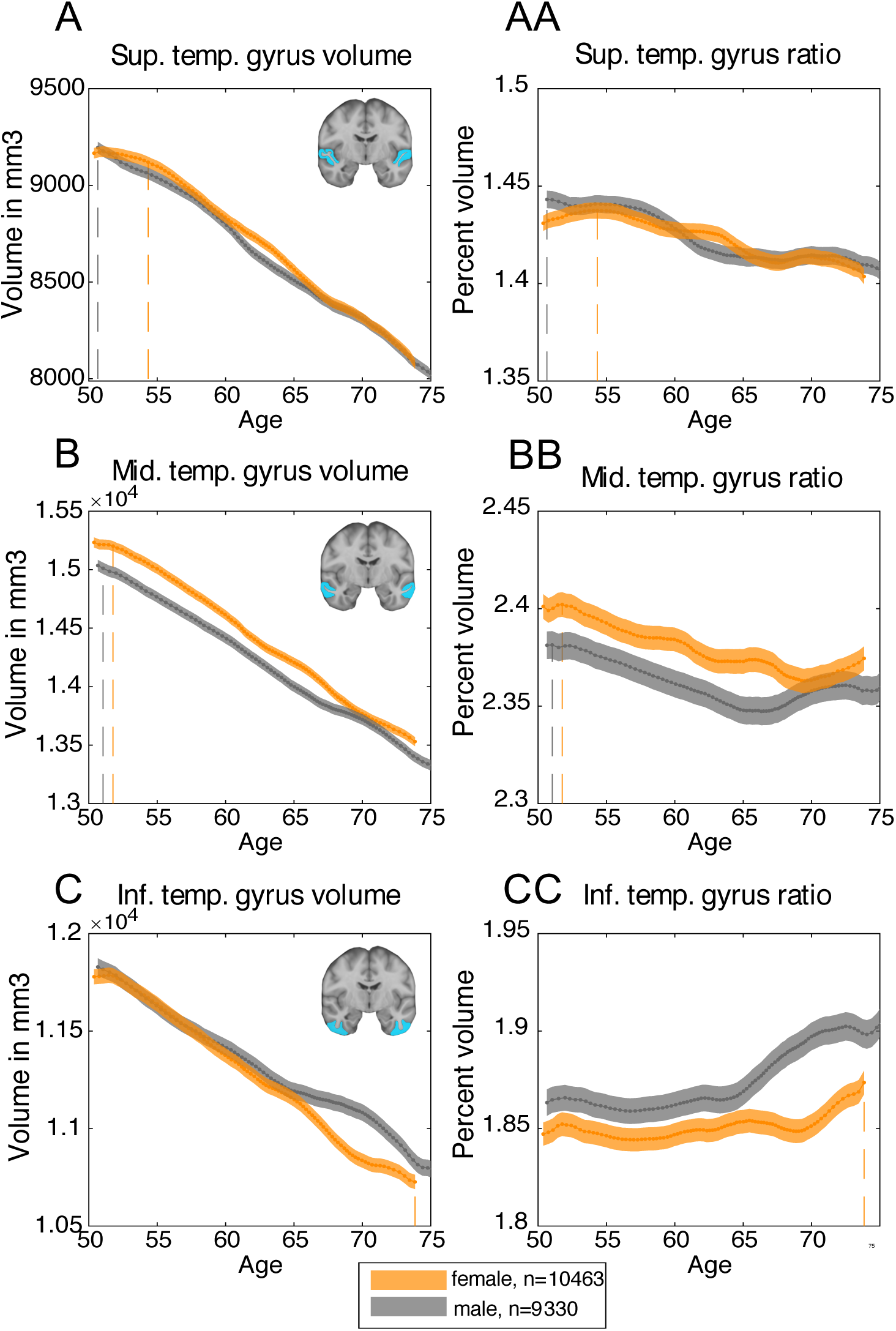
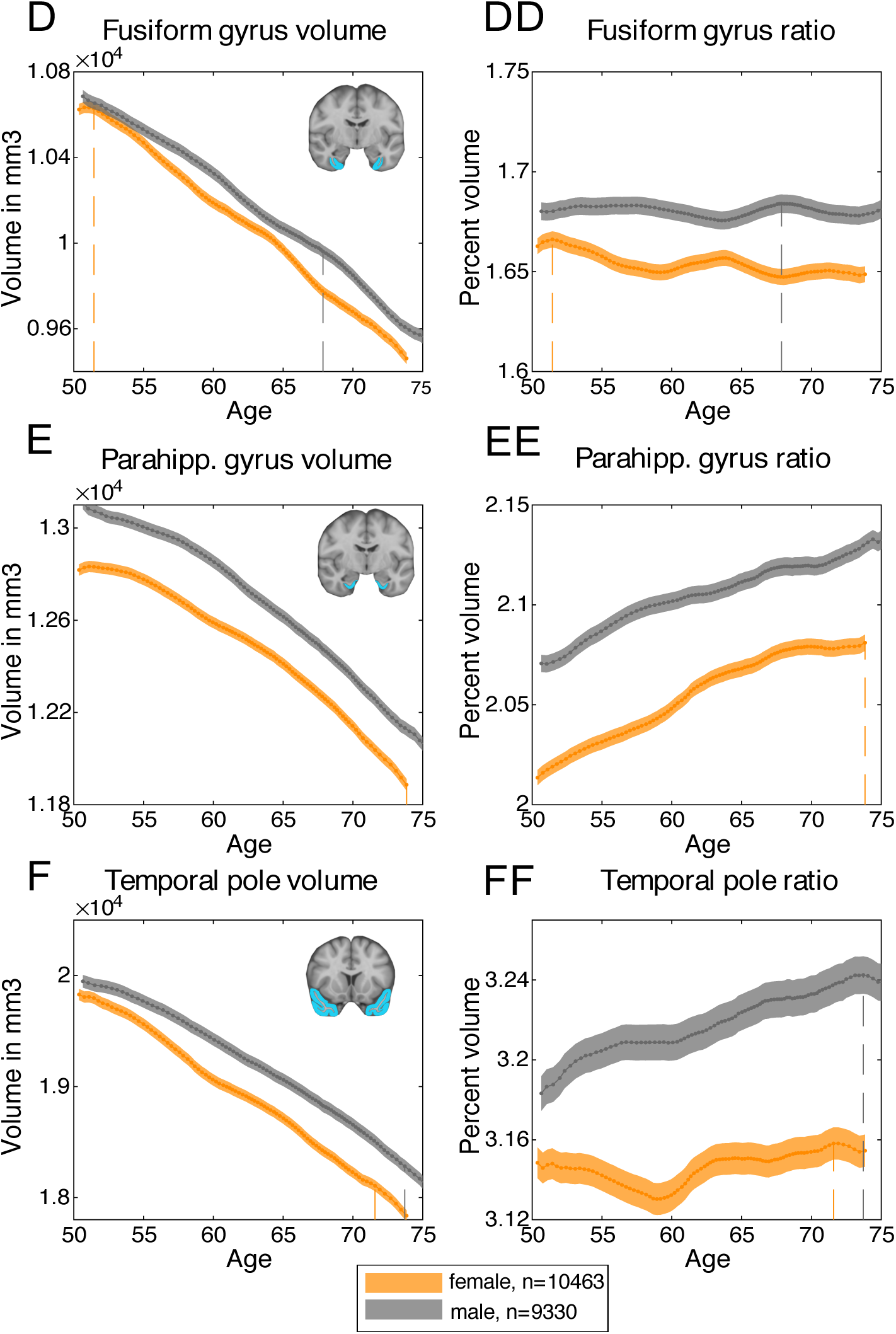
Trajectory of temporal lobe volumes with age, corrected for head-size. Dashed lines indicate points of maximum ratio. **A**. Bilateral superior temporal gyrus volume. **AA.** Bilateral superior temporal gyrus ratio to rest of grey matter. **B**. Bilateral middle temporal gyrus volume. **BB.** Bilateral middle temporal gyrus ratio to rest of grey matter. **C**. Bilateral inferior temporal gyrus volume. **CC.** Bilateral inferior temporal gyrus ratio to rest of grey matter. **D**. Bilateral fusiform gyrus volume. **DD.** Bilateral fusiform gyrus ratio to rest of grey matter. **E**. Bilateral parahippocampal gyrus volume. **EE.** Bilateral parahippocampal gyrus ratio to rest of grey matter. **F**. Bilateral temporal pole volume. **FF.** Bilateral temporal pole ratio to rest of grey matter.

In addition, the ages of the peak *ratios* with respect to the rest of grey matter differs between regions. For example, the ratios of superior (female: bin(*max ratio)* = 52.73 years – 55.85 years; male: bin(*max ratio*) = 44.45 years – 52.73 years) and middle temporal gyrus volume (female: bin(*max ratio)* = 49.30 years – 53.11 years; male: bin(*max ratio*) = 48.30 years – 53.11 years) to rest of grey matter peak in the youngest age groups of the sample. By contrast, ratios for inferior temporal gyrus (female: bin(*max ratio)* = 71.92 years – 80.65 years; male: bin(*max ratio*) = 73.41 years – 80.27 years), temporal pole (female: bin(*max ratio)* = 70.22 years – 73.45 years; male: bin(*max ratio*) = 72.25 years – 75.56 years) and parahippocampal volume (female: bin(*max ratio)* = 71.92 years – 80.65 years; male: bin(*max ratio*) = 73.41 years – 80.27 years) to the rest of the grey matter peak at the oldest age groups. The ratio of fusiform gyrus volume to the rest of grey matter volume does not show pronounced change over age. There were no significant differences in the age of peak ratio between males and females for any of these temporal lobe regions, calculated by permutation testing (*n(permutations)* = 5000*, all p* > .05), except for fusiform gyrus (*p* = .004).

## Discussion

Data from the UK Biobank Imaging were analysed to provide normative information on hippocampal volume as a function of age using the largest sample size published to date (*N*=19,793). We used a model-free sliding window approach to examine absolute and relative hippocampal volume, as well as neighbouring temporal lobe volumes, as a function of age. This approach was deemed most appropriate as it makes few assumptions, and does not impose a linear relationship between brain volume and age. The analysis resulted in normative reference percentiles – or nomograms – for total grey matter and hippocampal volume along the age distribution, facilitating assessment of an individual’s hippocampal volume in relation to their age-group (see our online tool at (http://www.smanohar.com/biobank/).

For comparison, it is noteworthy that estimated hippocampal volumes from previous studies on patients with MCI or AD fall far below the 50^th^ percentile in the healthy population presented here. For example, mean uncorrected hippocampal volume (bilateral) was 5.55 cm^3^ in an amnestic MCI group with an average age of 70.7 years (Vos et al. 2013). This would fall below the 2.5^th^ percentile in our sample. In a sample of controls and patients with MCI and AD of similar mean age as in our sample (67 years, 71 years, and 67 years, respectively), controls would score between our 50^th^ and 75^th^ percentiles, while patients score below the 5^th^ percentile (Henneman et al. 2009). Volumes compatible with our current results for healthy controls (3.57 cm^3^; mean age 75 years), but with far less dramatic reductions for MCI (3.26 cm^3^; mean age 73 years) and AD (2.97 cm^3^; mean age 72.6 years), were reported using data from the Alzheimer’s Disease Neuroimaging Initiative (ADNI) database (Mulder et al. 2014). These would likely correspond to healthy controls scoring around our 50^th^ percentile, but with patients scoring between the 5^th^ and 25^th^ percentiles.

It is likely that different methods, including sample selection as well as image acquisition, image processing, and hippocampal segmentation tool used, underlie these discrepancies. These issues highlight the importance of large datasets and unified analysis pipelines. Before efforts to harmonize analysis across MRI datasets solve some of these concerns, the nomograms presented here provide a robust comparator, when used with estimates obtained using the same, or similar, acquisition protocols, hardware and pre-processing pipeline.

It can also currently not easily be disambiguated whether a low percentile on the nomogram is due to pathological atrophy or is simply constitutional, e.g., due to an unusually small volume from birth. However, future research with longitudinal data from the UK Biobank may facilitate the resolution of some of these constraints.

Associations of hippocampal volume with education, smoking, and hypertension were significant, but effect sizes were small, and deconfounding for these variables did not effectively change any of the reported results. Since significant findings with small effect sizes are not necessarily practically relevant with big data, we decided not to correct for these factors in the calculations of percentiles in favour of generalisability. However, it should be noted that participants in UK Biobank appear to smoke less, have lower BMIs, and are generally healthier than would be expected in the general population of the United Kingdom (Fry et al. 2017).

The results of two different types of analysis revealed a slight acceleration of hippocampal volume loss around age 60-65 years for females, whereas for males the rate of hippocampal volume loss may increase earlier around 50 years (Figures 7 and 8). In addition, for both women and men, there was an increase in rate of hippocampal volume loss relative to the rest of the grey matter from around ages 67 and 63 years, respectively. Thus, hippocampal volume declined slower than the rest of grey matter until around age 67 in women, and age 63 in men, after which hippocampal volume declined faster than the rest of grey matter. This may indicate a particular vulnerability of the hippocampus in ageing, as this effect was specific to the hippocampus and was not found to this extent in neighbouring brain areas such as parahippocampal gyrus or temporal gyrus (Figure 9). While our method does not allow for precise age estimations, we show a robust age-range for the ratio peak with different window bins and smoothing kernels.

A previous cross-sectional study with results from 1100 participants reported a similar acceleration of rate of hippocampal volume loss (Fjell et al. 2013). Relationships of age and several brain volumes were analysed using a nonparametric smoothing spline approach. The authors reported an acceleration of rate of hippocampal volume loss starting around 50 years of age, with a marked increase in rate of hippocampal volume loss around 60 years. In agreement with our findings, no such acceleration point was found in grey matter. However, in this study no sex differences in these curves were calculated. In contrast, other cross-sectional studies using linear regression to estimate associations of age with hippocampal volume found an acceleration of volume loss at a slightly later age of 72 years (Zhang et al. 2010), or no acceleration point (Knoops et al. 2012).

In the analyses presented here, the significant slope changes at the estimated joinpoints suggest, however, that a linear fit is not the most appropriate approach to measure the association between age and relative hippocampal volume in our sample. Importantly, the trajectories of other temporal lobe volumes, as well as the rest of grey matter across age do not show the same acceleration of volume loss as observed in hippocampal volume. This may underlie the specific vulnerability of hippocampal volume loss in older age. Yet, the findings presented here vary slightly depending on method used (e.g. sliding window or joinpoint), and need to be validated with longitudinal rather than cross-sectional data. For example, the current data do not allow for conclusions of whether those with the fastest decline, or those with the lowest baseline volume, are at more risk of developing dementia. With the availability of longitudinal health outcomes, future studies may also explore the role of the dementia-associated APOE ε4 allele on hippocampal volume across age.

In the absence of differential age-effects on grey matter volume between males and females, we found significantly larger mean total grey matter, and middle temporal gyrus volumes in females after correcting for head size and age. However, effect sizes were small. On the other hand, males had moderately larger parahippocampal and temporal pole volumes. The data presented here also indicate hemispheric asymmetry of the tested brain volumes. The right hippocampus was slightly, but significantly, larger than the left hippocampus for both men and women. Right-larger-than-left asymmetry was also found in the superior and inferior temporal gyri. In contrast, there was considerable left-larger-than-right asymmetry for both the parahippocampal and fusiform gyrus. The observed asymmetries, and the differences in asymmetry direction, may be the result of noise in the MRI signals and imperfections of the automated volume segmentation and quality control tools used here. For example, the algorithm that the segmentation uses to determine which voxel is classified as ‘parahippocampal’ versus ‘hippocampal’ may be the reason for the differences in asymmetry direction between the two volumes. Thus, while these results suggest some structural differences between hemispheres, such as asymmetry of convolutions, it is unclear what these differences entail precisely. However, the right-larger-than-left asymmetry of the hippocampus has been reported previously in smaller cohorts of younger and older adults (Zhang et al. 2010; Wellington et al. 2013; Pedraza et al. 2018), as well as in patients with AD and MCI (Shi et al. 2009). A recent study reporting on 400 participants from the ADNI dataset also found increasing right-larger-than-left hippocampus asymmetry going from healthy controls, to patients with MCI, to patients with AD (Alberich-Bayarri et al. 2018). As the left hippocampus is smaller even in healthy ageing, the increasing asymmetry in the course of AD may be explained by greater vulnerability to pathology of the left compared to the right hippocampus. What is less clear is how the left-larger-than-right asymmetries of the parahippocampal and fusiform gyri may be affected by ageing and AD pathology.

Functional lateralisation has been shown for a number of brain processes, such as language (Vigneau et al. 2006), face perception (Zhen et al. 2015), or visual processing (Zhen et al. 2017), which may explain the asymmetries observed here in other areas such as the inferior and superior temporal gyri. However, the precise nature of the relationship between brain structural lateralisation and function remains unclear (Bishop 2013; Batista-García-Ramó and Fernández-Verdecia 2018), highlighting the importance of considering confounding factors for the analysis of norm values. We therefore provide separate nomograms for left and right hippocampus, as well as for males and females.

## Conclusion

Analysis of 19,793 generally healthy participants in the UK Biobank revealed effects of age, sex, and hemisphere on selected temporal lobe volumes. The data provide normative values for hippocampal and total grey matter volume as a function of age for reference in research and clinical settings, based on an unprecedented sample size. These norm values may be used together with automated percentile estimation tools such as the one described here to provide a rapid, but objective, evaluation of the patient’s hippocampal volume status. While the current findings are based on cross-sectional data, the longitudinal health outcomes that will become available in UK Biobank over the next years will add invaluable information to the role of hippocampal atrophy in ageing.

## Acknowledgements

### Funding

This research was conducted using the UK Biobank resource (application ID: 32011). UK Biobank was established by the Wellcome Trust, the Medical Research Council, the United Kingdom Department of Health, and the Scottish Government. The UK Biobank has also received funding from the Welsh Assembly Government, the British Heart Foundation, and Diabetes UK.

This work was supported by the Wellcome Trust (206330/Z/17/Z to M.H.), the Medical Research Council (DPhil Studentship to L.N and a Clinician Scientist Fellowship to SGM). SMS, FA-A, MJ, and CEM acknowledge support from the National Institute for Health Research (NIHR) Oxford Health Biomedical Research Centre (BRC), a partnership between Oxford Health NHS Foundation Trust and the University of Oxford. The Wellcome Centre for Integrative Neuroimaging is supported by core funding from the Wellcome Trust (203139/Z/16/Z).

