## Supplementary Material for "Hippocampal volume across age: Nomograms derived from over 19,700 people in UK Biobank"

**Suppl. Table S1**: Brain volume by hypertension status and BMI correlations
**Suppl. Table S2**: Brain volume by education level
**Suppl. Table S3:** Brain volume by smoking status
**Suppl. Table S4**: Brain volume by hemisphere
**Suppl. Table S5**: Brain volume by sex
**Suppl. Table S6:** Model estimates from joinpoint regression of total grey matter

**Suppl. Figure S1:** Difference in brain volume by hypertension
**Suppl. Figure S2**: Difference in brain volume by education level
**Suppl. Figure S3**: Difference in brain volume by smoking status

**Suppl. Figure S4**: Nomogram of head size corrected left hippocampus for females
**Suppl. Figure S5**: Nomogram of head size corrected left hippocampus for males
**Suppl. Figure S6:** Nomogram of head size corrected right hippocampus for females
**Suppl. Figure S7**: Nomogram of head size corrected right hippocampus for males
**Suppl. Figure S8:** Nomogram of uncorrected total grey matter for females
**Suppl. Figure S9:** Nomogram of uncorrected total grey matter for males
**Suppl. Figure S10:** Nomogram of head size corrected total grey matter for females
**Suppl. Figure S11**: Nomogram of head size corrected total grey matter for males

**Suppl. Figure S12:** Sliding-window curves with fixed age-bins for head size corrected hippocampus
**Suppl. Figure S13**: Sliding-window curves for left and right hippocampal volume across age
**Suppl. Figure S14:** Slope of hippocampal volume by age
**Suppl. Figure S15:** Sliding-window curves (Fig.7) without smoothing
**Suppl. Figure S16:** Sliding-window curves Fig.7) with smoothing kernel 10
**Suppl. Figure S17:** Sliding-window curves (Fig.7) with 20% quantile width and smoothing kernel 20
**Suppl. Figure S18**: Sliding-window curves (Fig.7) with 20% quantile width and smoothing kernel 10

**Suppl. Table 1:** *Brain volume by hypertension status and BMI correlations**

| | **No Hypertension** *(Mean ± std)* | **Hypertension** *(Mean ± std)* | ***p*-value**, Effect size** | **BMI** *(Pearson's r, p-value**)* |
|---|---|---|---|---|
| **Total grey matter volume** | 617,487.42 ± 26,298.42 $mm^3$ | 612,994.27 ± 27938.30 $mm^3$ | *p* < .001, *Hedges' G* = .16 | *r* = -.11, *p* < .001 |
| **Hippocampal volume** | 3868.40 ± 351.77 $mm^3$ | 3846.02 ± 359.61 $mm^3$ | *p* < .001, *Hedges' G* = .07 | *r* = -.03, *p* < .001 |
| **Parahippocampal gyrus** | 8440.67 ± 591.44 $mm^3$ | 8396.09 ± 605.95 $mm^3$ | *p* < .001, *Hedges' G* = .08 | *r* = -.04, *p* < .001 |
| **Fusiform gyrus** | 7799.46 ± 719.50 $mm^3$ | 7747.10 ± 715.59 $mm^3$ | *p* < .001, *Hedges' G* = .07 | *r* = -.05, *p* < .001 |
| **Inferior temporal gyrus** | 8431.44 ± 1033.65 $mm^3$ | 8394.17 ± 1018.81 $mm^3$ | *p* = .03 *Hedges' G* = .04 | *r* = -.02, *p* = .009 |
| **Mid temporal gyrus** | 10,744.31 ± 1110.08 $mm^3$ | 10,602.11 ± 1109.54 $mm^3$ | *p* < .001, *Hedges' G* = .13 | *r* = -.04, *p* < .001 |
| **Superior temporal gyrus** | 6364.29 ± 709.80 $mm^3$ | 6297.34 ± 710.23 $mm^3$ | *p* < .001, *Hedges' G* = .09 | *r* = -.03, *p* = .001 |
| **Temporal pole** | 9507.25 ± 892.26 $mm^3$ | 9460.28 ± 918.43 $mm^3$ | *p* = .002 *Hedges' G* = .05 | *r* = -.02, *p* = .001 |
| **N (Hippocampus)** | 15,367 | 4426 | / | / |

** Volumes are averaged over hemisphere and corrected for head-size and age*
*** Bonferroni-corrected $\alpha$ = .05/8 multiple comparisons = .006*

**Suppl. Table 2:** *Brain volume by education level**

| | College/University (*Mean ± std*) | Non-university (*Mean ± std*) | *p*-value**, Effect size |
|---|---|---|---|
| **Total grey matter volume** | 616,558.75 ± 26,825.60 $mm^3$ | 616,444.71 ± 26,528.94 $mm^3$ | $p = .77$ |
| **Hippocampal volume** | 3872.48 ± 358.24 $mm^3$ | 3848.42 ± 345.29 $mm^3$ | $p < .001$, *Hedges' G* = .07 |
| **Parahippocampal gyrus** | 8436.36 ± 596.68 $mm^3$ | 8422.77 ± 591.79 $mm^3$ | $p = .12$ |
| **Fusiform gyrus** | 7801.96 ± 729.04 $mm^3$ | 7766.99 ± 701.35 $mm^3$ | $p = .001$, *Hedges' G* = .05 |
| **Inferior temporal gyrus** | 8433.30 ± 1040.77 $mm^3$ | 8410.89 ± 1012.63 $mm^3$ | $p = .14$ |
| **Mid temporal gyrus** | 10,705.05 ± 1118.75 $mm^3$ | 10,728.99 ± 1109.93 $mm^3$ | $p = .14$ |
| **Superior temporal gyrus** | 6350.49 ± 717.46 $mm^3$ | 6349.16 ± 701.52 $mm^3$ | $p = .89$ |
| **Temporal pole** | 9514.02 ± 906.29 $mm^3$ | 9468.41 ± 884.33 $mm^3$ | *p* < 001, *Hedges' G* = .05 |
| **N (Hippocampus)** | 12275 | 7371 | / |

** Volumes are averaged over hemisphere and corrected for head-size and age*
*** Bonferroni-corrected $\alpha$ = .05/8 multiple comparisons = .006*

**Suppl. Table 3:** *Brain volume by smoking status**

**A**

| | **Never Smoked** *(Mean ± std)* | **Previous Smoker** *(Mean ± std)* | **Current Smoker** *(Mean ± std)* | ***F-test**** |
|---|---|---|---|---|
| **Total grey matter volume** | 618,402.92 ± 26,445.67 $mm^3$ | 613,808.99 ± 26,817.75 $mm^3$ | 609,778.46 ± 27,280.52 $mm^3$ | $F(3,19779) = 60.12$, $p < .001$ |
| **Hippocampal volume** | 3871.85 ± 353.02 $mm^3$ | 3853.36 ± 353.75 $mm^3$ | 3819.70 ± 353.53 $mm^3$ | $F(3,19541) = 8.26$, $p < .001$ |
| **Parahippocampal gyrus** | 8444.84 ± 592.86 $mm^3$ | 8414.13 ± 594.62 $mm^3$ | 8359.77 ± 617.07 $mm^3$ | $F(3,19660) = 7.65$, $p < .001$ |
| **Fusiform gyrus** | 7808.68 ± 717.71 $mm^3$ | 7759.14 ± 719.81 $mm^3$ | 7732.03 ± 720.93 $mm^3$ | $F(3,19660) = 8.62$, $p < .001$, |
| **Inferior temporal gyrus** | 8445.65 ± 1030.63 $mm^3$ | 8402.03 ± 1025.52 $mm^3$ | 8297.72 ± 1051.02 $mm^3$ | $F(3,19568) = 6.81$, $p < .001$ |
| **Mid temporal gyrus** | 107,46.47 ± 1109.89 $mm^3$ | 10,668.04 ± 1104.04 $mm^3$ | 10,583.20 ± 1185.32 $mm^3$ | $F(3,19609) = 10.98$, $p < .001$ |
| **Superior temporal gyrus** | 6371.91 ± 714.98 $mm^3$ | 6317.67 ± 702.42 $mm^3$ | 6279.76 ± 705.16 $mm^3$ | $F(3,19616) = 10.91$, $p < .001$ |
| **Temporal pole** | 9527.57 ± 899.46 $mm^3$ | 9453.23 ± 891.44 $mm^3$ | 9382.81 ± 923.98 $mm^3$ | $F(3,19719) = 14.11$, $p < .001$ |
| **N (Hippocampus)** | 122232 | 6597 | 759 | / |

** Volumes are averaged over hemisphere and corrected for head-size and age*
*** Bonferroni-corrected $\alpha$ = .05/8 multiple comparisons = .006*

**B**

| | Never Smoked VS Previous Smoker | Previous Smoker VS Current Smoker | Never smoked VS Current Smoker |
|---|---|---|---|
| **Total grey matter volume** | ↑ | ↑ | ↑ |
| **Hippocampal volume** | ↑ | ↑ | ↑ |
| **Parahippocampal gyrus** | No difference | ↑ | ↑ |
| **Fusiform gyrus** | ↑ | No difference | ↑ |
| **Inferior temporal gyrus** | No difference | ↑ | ↑ |
| **Mid temporal gyrus** | ↑ | No difference | ↑ |
| **Superior temporal gyrus** | ↑ | No difference | ↑ |
| **Temporal pole** | ↑ | No difference | ↑ |

**↑ = significantly larger volume**

**Suppl. Table 4:** *Brain volume by hemisphere*

| | **Left hemisphere** *(Mean ± std)* | **Right hemisphere** *(Mean ± std)* | ***p*-value*, Effect size** |
|---|---|---|---|
| **Hippocampal volume** | 3807.70 ± 451.63 $mm^3$ | 3915.91 ± 464.43 $mm^3$ | $p < .001$, *Hedges' G* = .24 |
| **Parahippocampal gyrus** | 4642.01 ± 537.72 $mm^3$ | 4383.59 ± 551.59 $mm^3$ | $p < .001$, *Hedges' G* = .48 |
| **Fusiform gyrus** | 5477.83 ± 713.48 $mm^3$ | 4621.58 ± 612.06 $mm^3$ | $p < .001$, *Hedges' G* = 1.30 |
| **Inferior temporal gyrus** | 5580.50 ± 942.11 $mm^3$ | 5689.91 ± 904.90 $mm^3$ | $p < .001$, *Hedges' G* = .12 |
| **Middle temporal gyrus** | 7155.30 ± 1076.31 $mm^3$ | 7121.20 ± 1017.23 $mm^3$ | $p = .61$ |
| **Superior temporal gyrus** | 4055.07 ± 647.87 $mm^3$ | 4591.15 ± 694.65 $mm^3$ | $p < .001$, *Hedges' G* = .80 |
| **Temporal pole** | 9523.30 ± 1260.67 $mm^3$ | 9473.84 ± 1213.86 $mm^3$ | $p <.001$, *Hedges' G* = .04 |

** Bonferroni-corrected $\alpha$ = .05/7 multiple comparisons = .007*

**Suppl. Table 5:** *Brain volume by sex*

| | **Male** (*Mean ± std*) | **Female** (*Mean ± std*) | ***p*-value**, Effect size** |
|---|---|---|---|
| **Total grey matter volume** | 614,423.67 ± 28,265.18 $mm^3$ | 618,328.53 ± 25,151.14 $mm^3$ | $p < .001$, *Hedges' G* = .15 |
| **Hippocampal volume** | 3866.54 ± 383.33 $mm^3$ | 3860.62 ± 325.01 $mm^3$ | $p = .24$ |
| **Parahippocampal gyrus** | 8499.62 ± 619.13 $mm^3$ | 8369.57 ± 565.75 $mm^3$ | $p < .001$, *Hedges' G* = .22 |
| **Fusiform gyrus** | 7825.78 ± 749.22 $mm^3$ | 7754.93 ± 689.34 $mm^3$ | $p < .001$, *Hedges' G* = .10 |
| **Inferior temporal gyrus** | 8454.76 ± 1076.24 $mm^3$ | 8395.22 ± 987.53 $mm^3$ | $p < .001$, *Hedges' G* = .06 |
| **Middle temporal gyrus** | 10,643.07 ± 1152.80 $mm^3$ | 10,774.03 ± 1069.91 $mm^3$ | $p < .001$, *Hedges' G* = .12 |
| **Superior temporal gyrus** | 6331.83 ± 732.669 $mm^3$ | 6364.79 ± 689.78 $mm^3$ | $p = .001$, *Hedges' G* = .05 |
| **Temporal pole** | 9571.71 ± 941.40 $mm^3$ | 9429.92 ± 852.70 $mm^3$ | $p < .001$, *Hedges' G* = .16 |

** Volumes are averaged over hemisphere and corrected for head-size and age*

*** Bonferroni-corrected $\alpha$ = .05/8 multiple comparisons = .006*

**Suppl. Table 6:** *Model Estimates from joinpoint regression of total grey matter*

| | Female | Male |
|---|---|---|
| **DF** | 21 | 21 |
| **Joinpoint** | 56-57 years | 62-63 years |
| **Joinpoint 95% LCL** | 53-54 years | 57-58 years |
| **Joinpoint 95% UCL** | 58-59 years | 66-67 years |
| **Slope Change Estimate** | -1273.98 $mm^3$ | -754.14 $mm^3$ |
| **Slope Change Std Error** | 427.19 $mm^3$ | 154.85 $mm^3$ |
| **Slope Change Test Statistic** | -2.98 | -4.87 |
| **Slope Change p-value** | .007 | < .0001 |

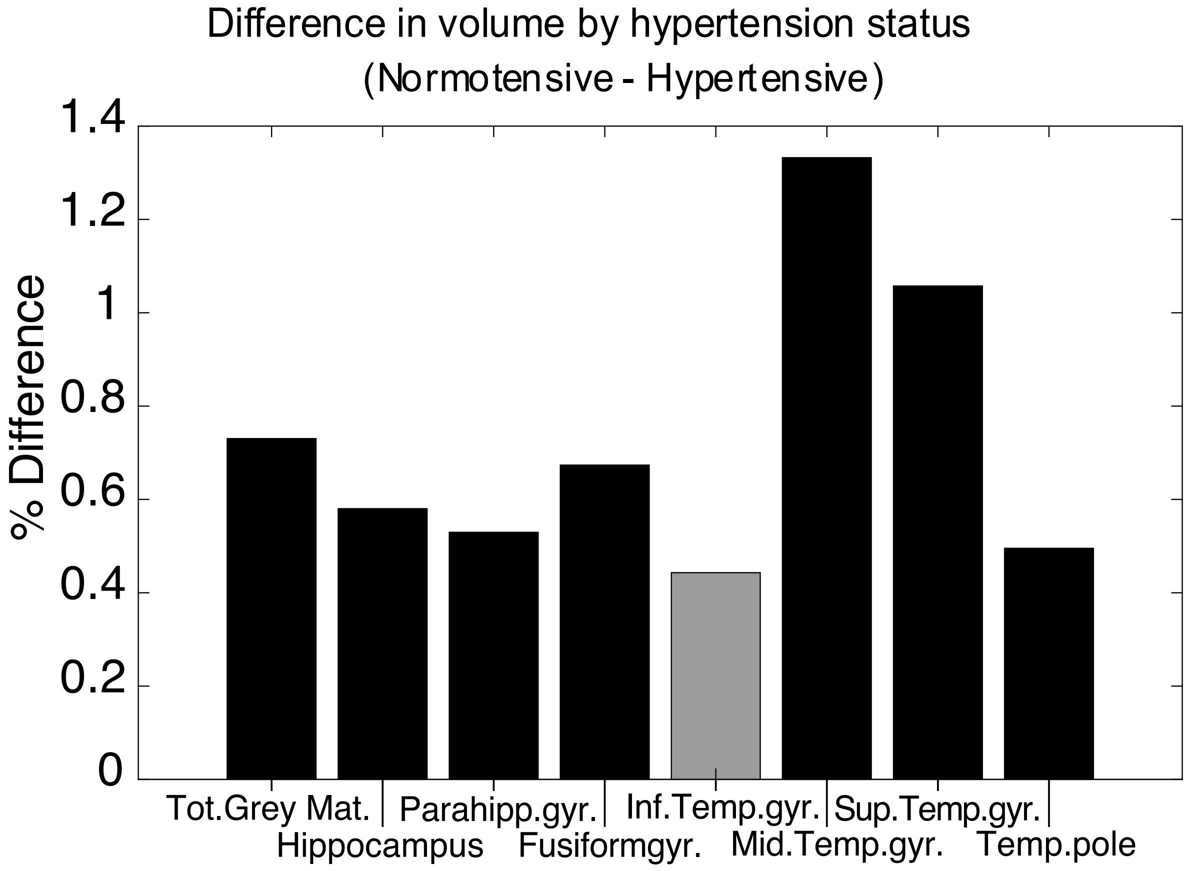


**Suppl. Figure 1:** *Difference in brain volume by hypertension status*

Percent difference in volumes was calculated with $\frac{|normotensive-hypertensive|}{\frac{normotensive+hypertensive}{2}} * 100.$ Positive percent differences correspond to larger volumes in participants without hypertension. Black bars indicate a significant corresponding t-test for volume differences at *p < .005 (Bonferroni-corrected* $\alpha$ *= .05/8 = .006)*

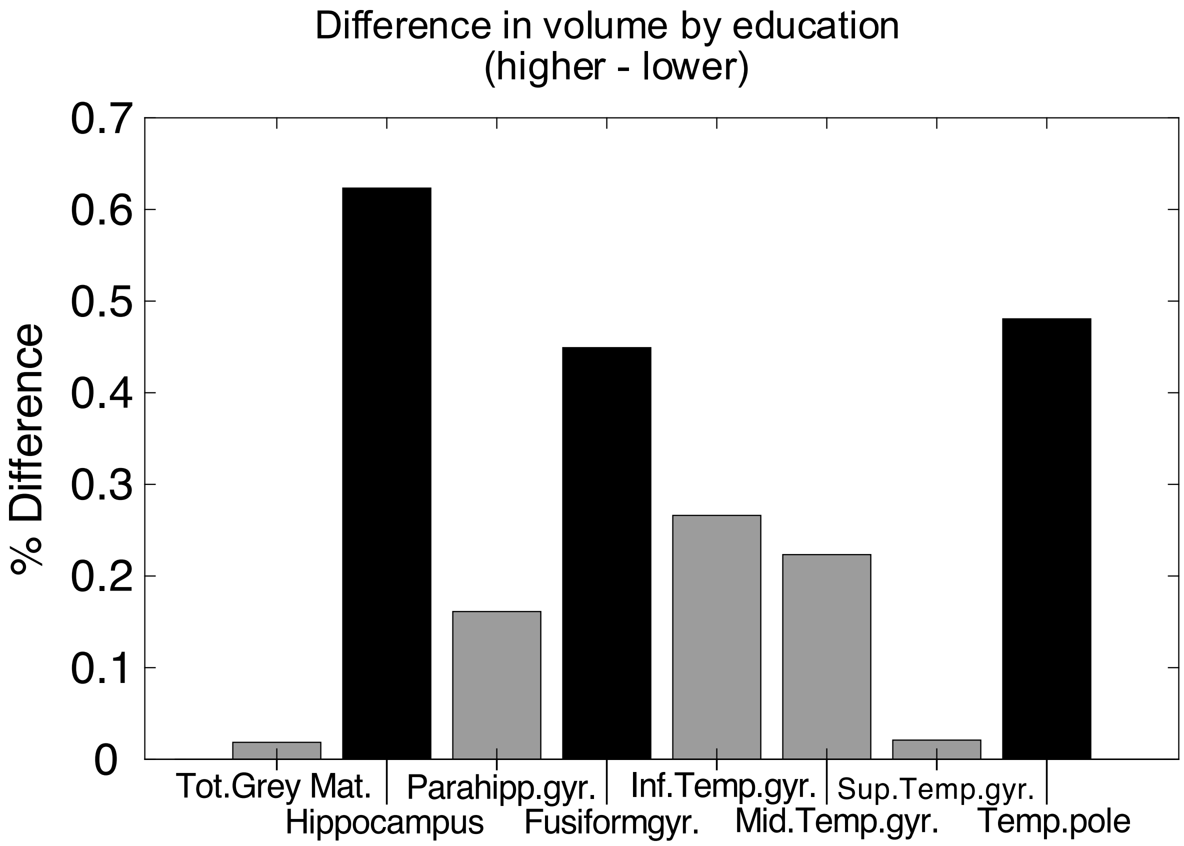


**Suppl. Figure 2:** *Percent volume differences between participants with lower and higher education levels*

Percent difference in volumes was calculated with $\frac{|higher-lower|}{\frac{higher+lower}{2}} * 100$. Positive percent differences correspond to larger volumes in participants with higher education levels. Black bars indicate a significant corresponding t-test for volume differences at *p < .001 (Bonferroni-corrected* $\alpha$ *= .05/8 = .006).*

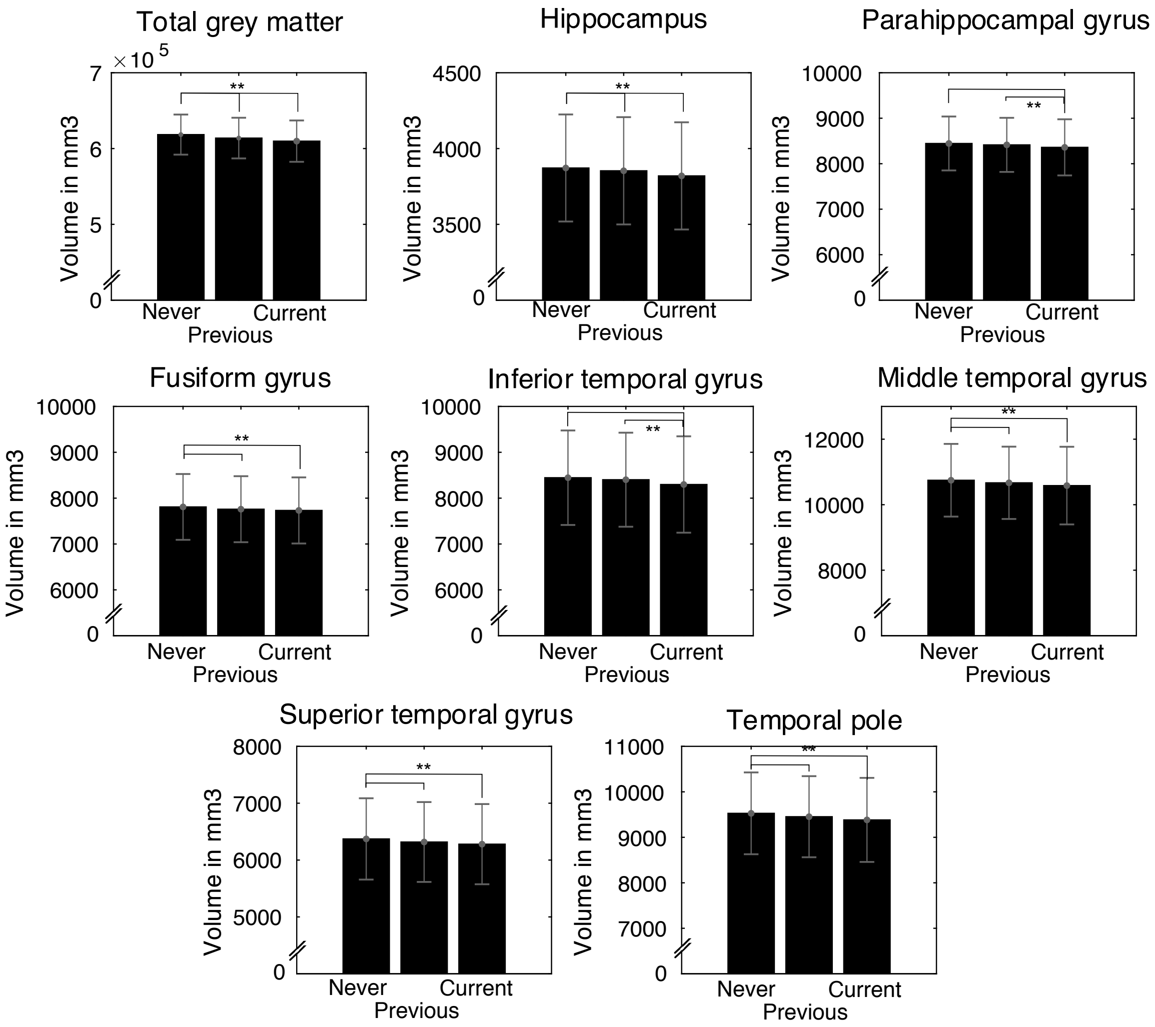


**Suppl. Figure 3***: Bar graphs for brain volume by smoking status.*

Participants indicated whether they are current smokers, previous smokers, or whether they have never smoked. Error bars show standard deviation.

**significant at $p < .001$ *(Bonferroni-corrected* $\alpha = .05/8 = .006$*)*

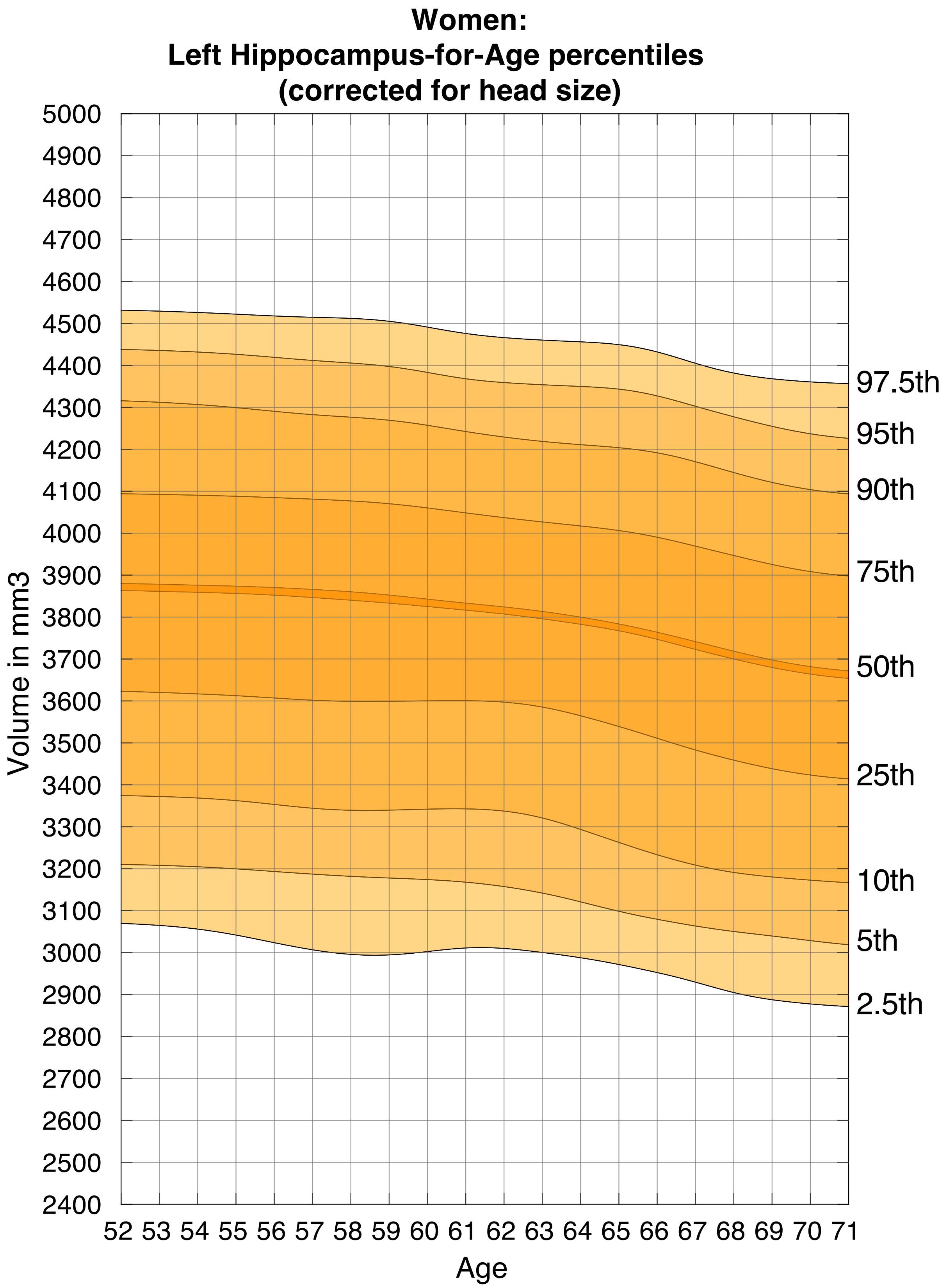


**Suppl. Figure S4**: *Nomogram of head size corrected left hippocampus for females*

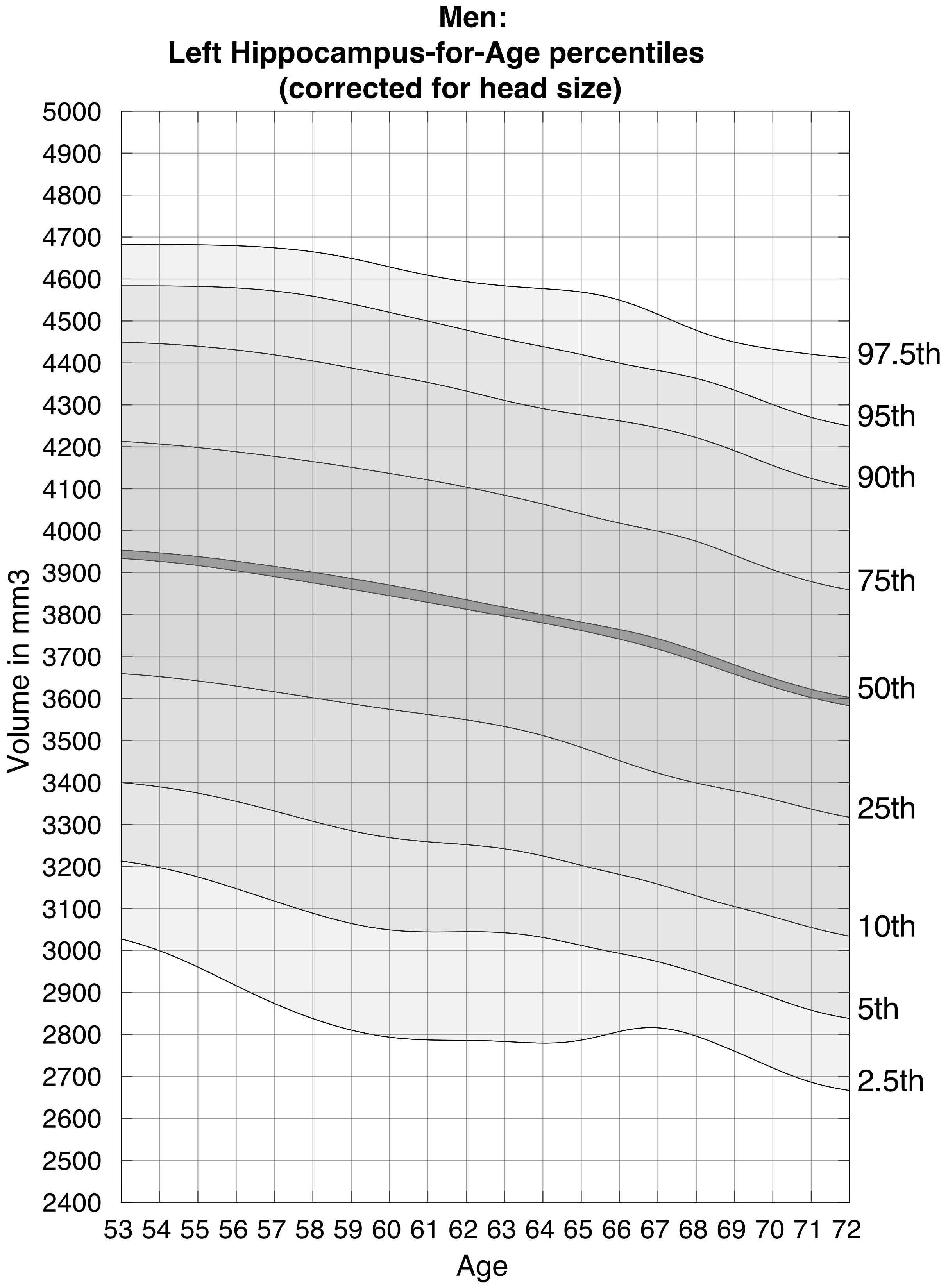


**Suppl. Figure S5**: *Nomogram of head size corrected left hippocampus for males*

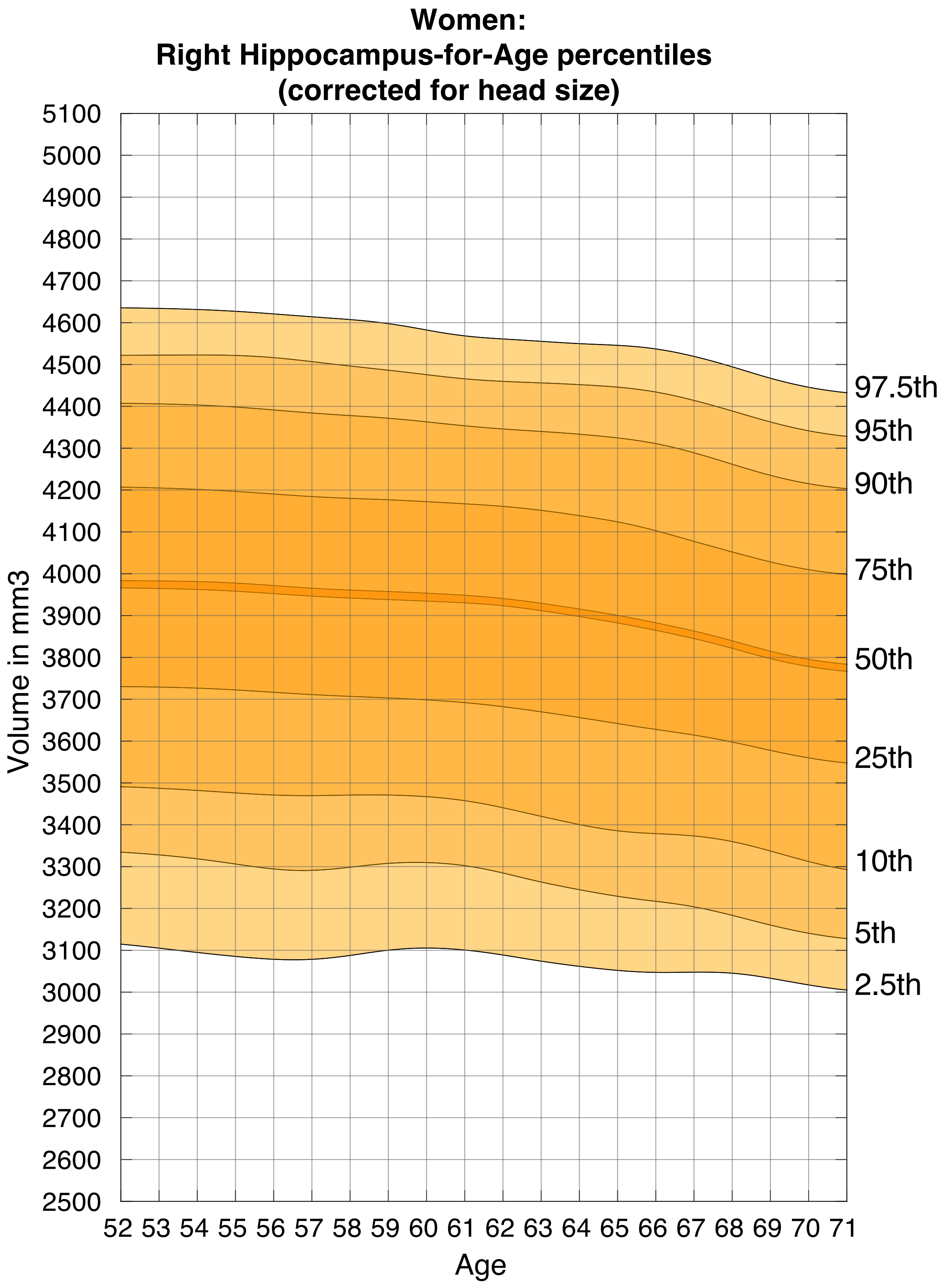


**Suppl. Figure S6***: Nomogram of head size corrected right hippocampus for females*

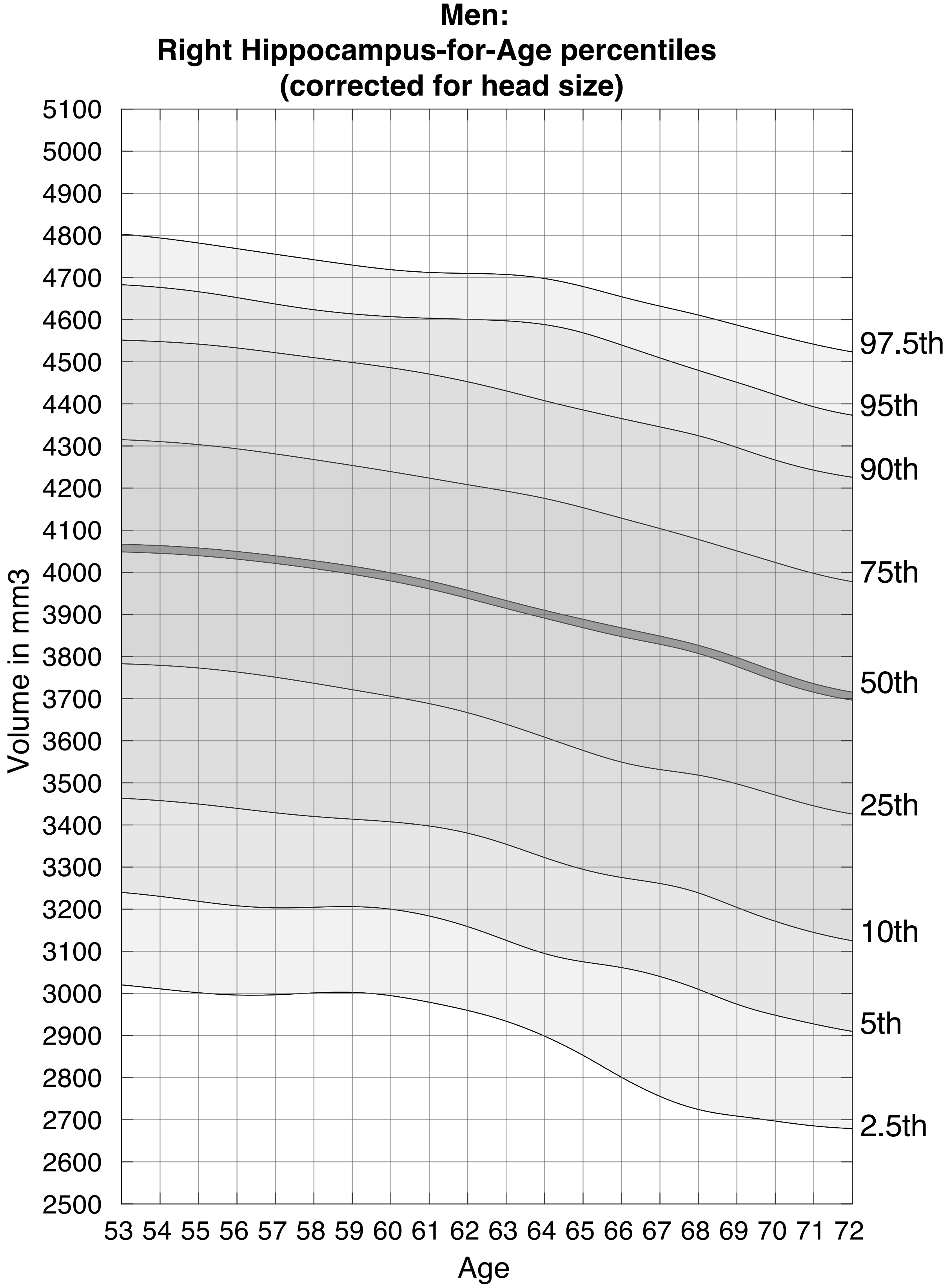


**Suppl. Figure S7**: *Nomogram of head size corrected right hippocampus for males*

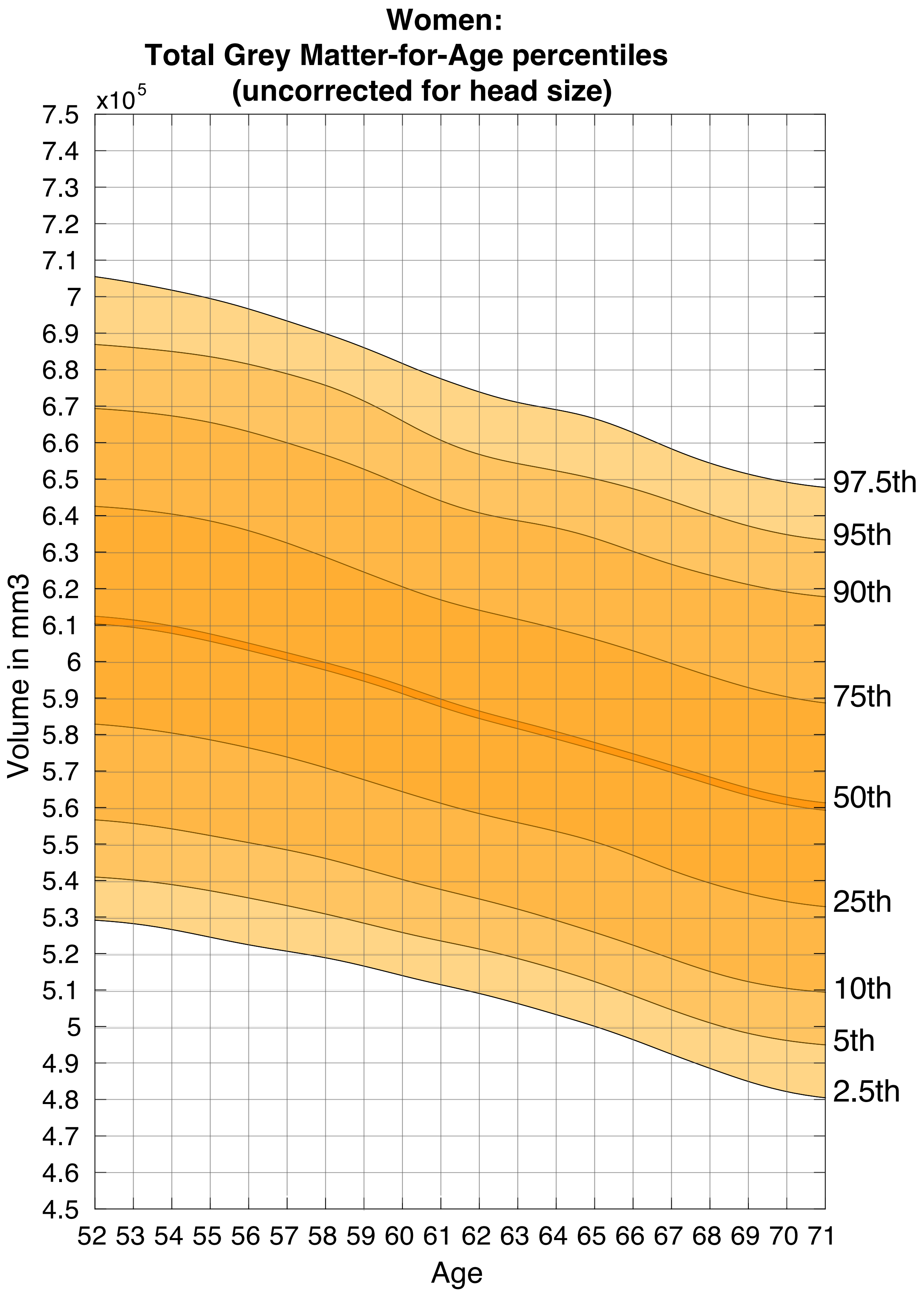


**Suppl. Figure S8:** *Nomogram of head size un-corrected total grey matter for females*

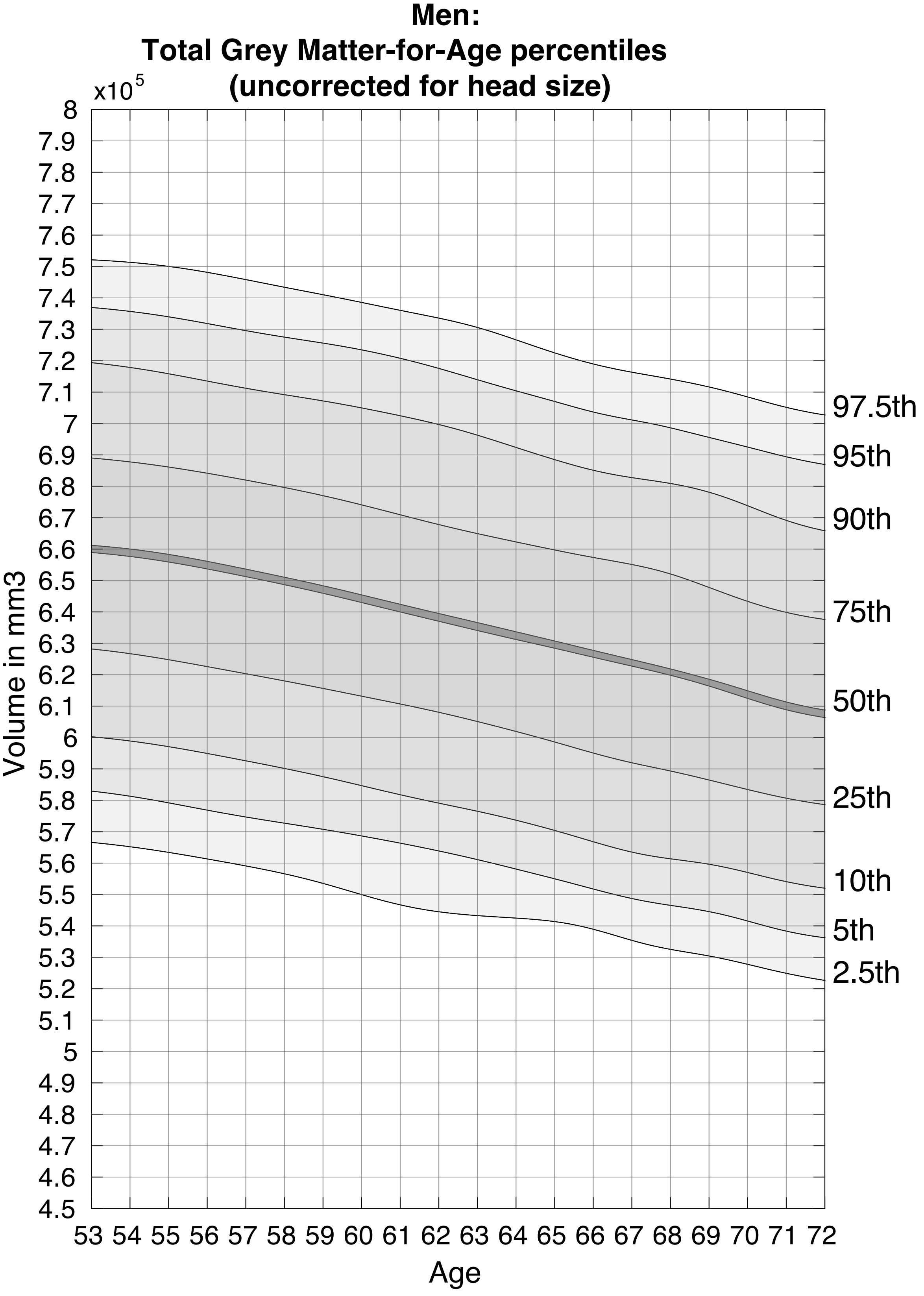


**Suppl. Figure S9***: Nomogram of head size un-corrected total grey matter for females*

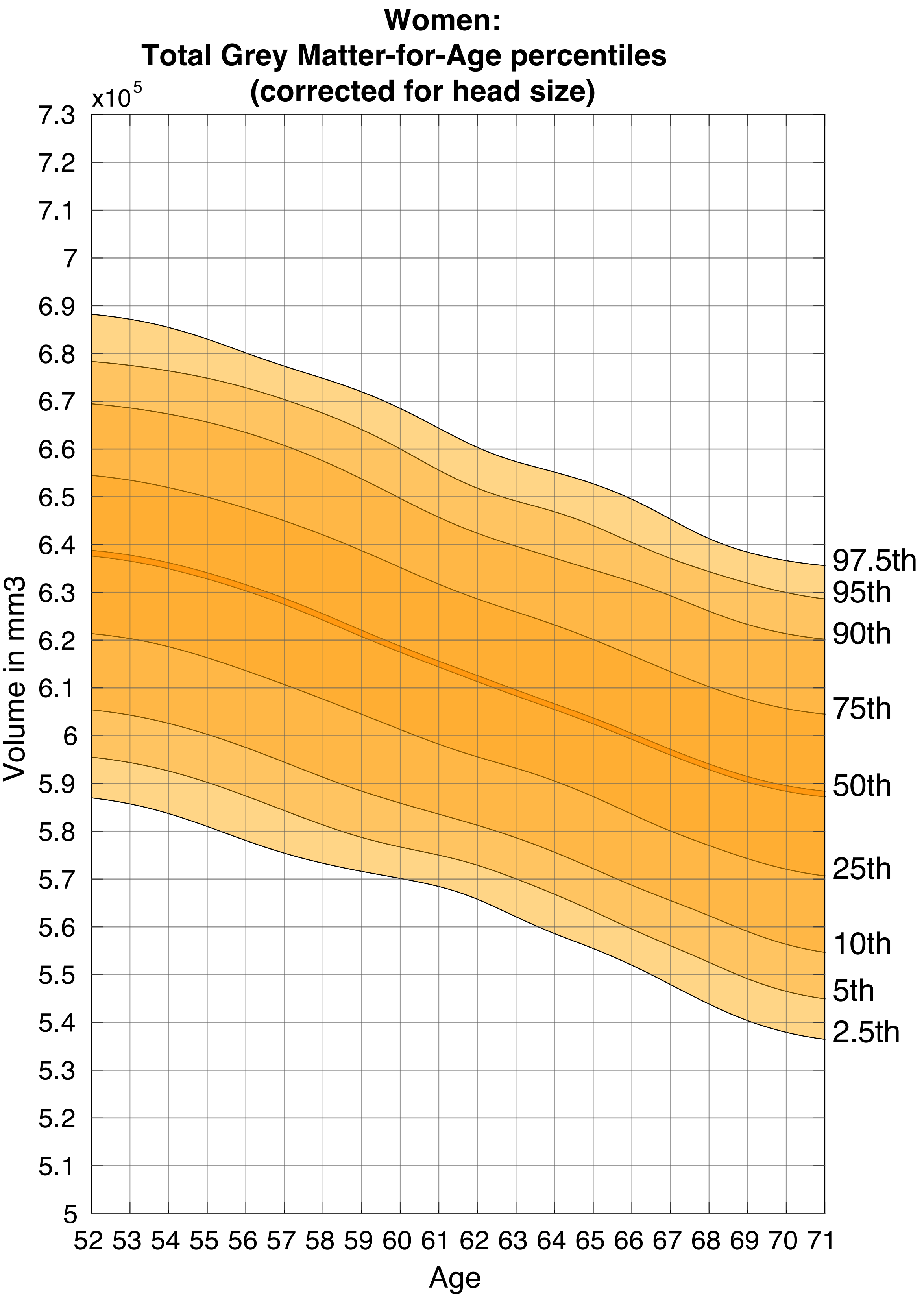


**Suppl. Figure S10:** *Nomogram of head size corrected total grey matter for females*

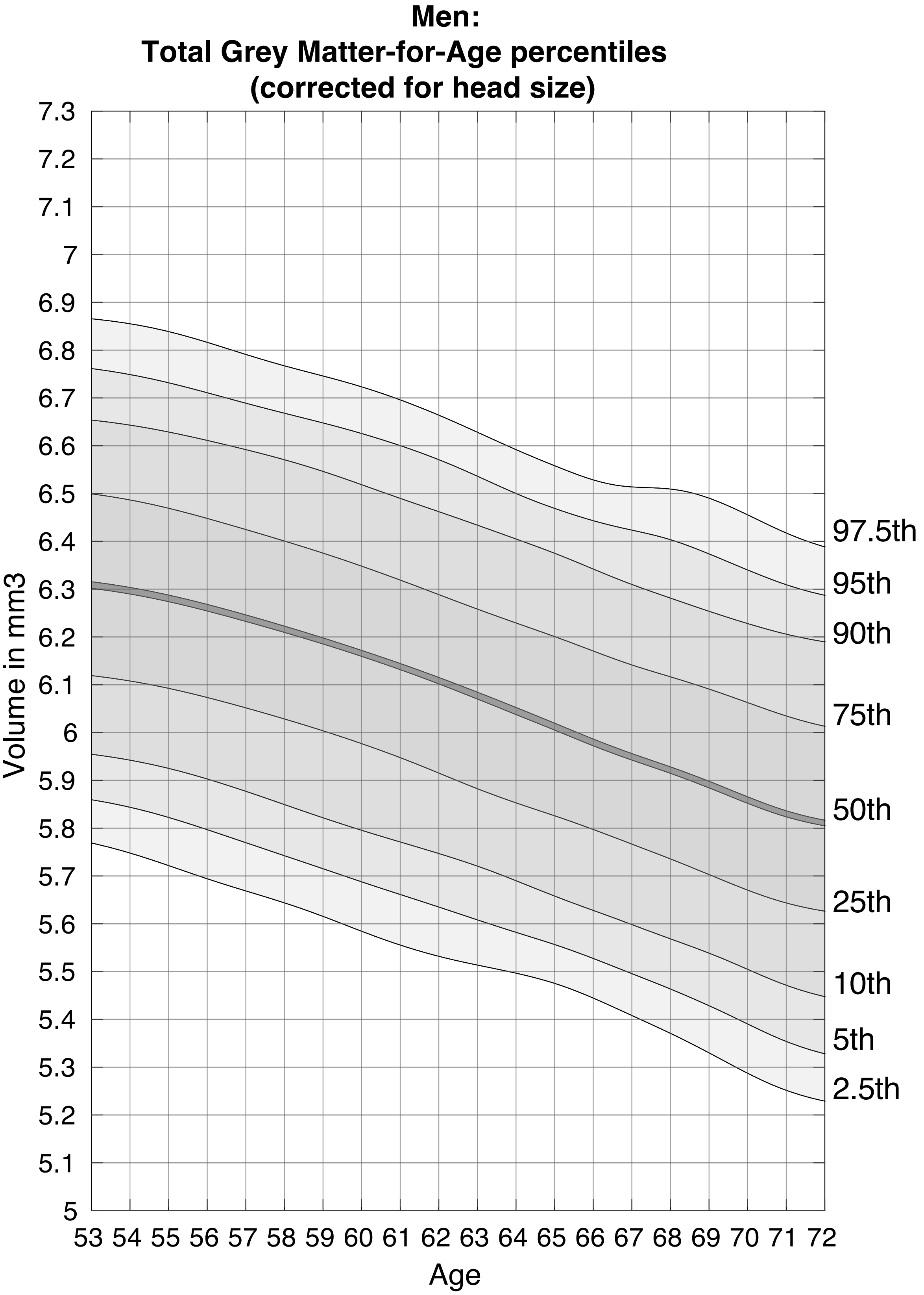


**Suppl. Figure S11**: *Nomogram of head size corrected total grey matter for males*

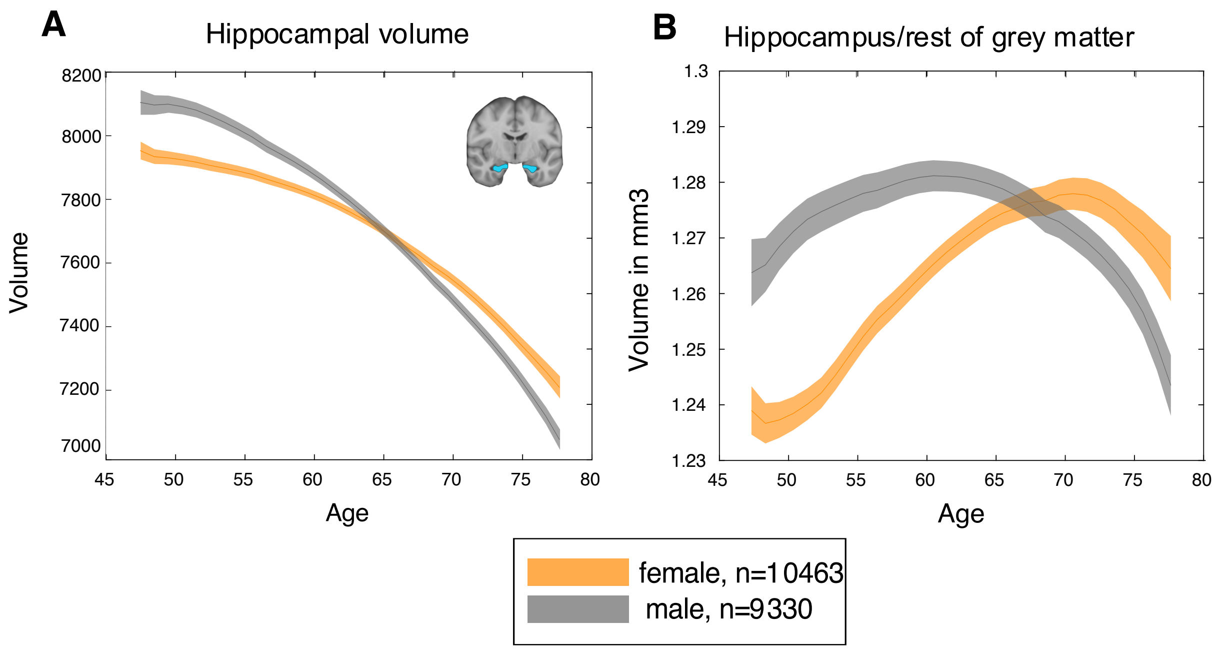


**Suppl. Figure S12:** *Sliding-window curves with fixed age-bins for head size corrected hippocampus*

**A**. Mean bilateral hippocampal volume including standard errors as a function of age, corrected for head size. **B**. Mean hippocampal volume to rest of grey matter ratio including standard errors as a function of age.

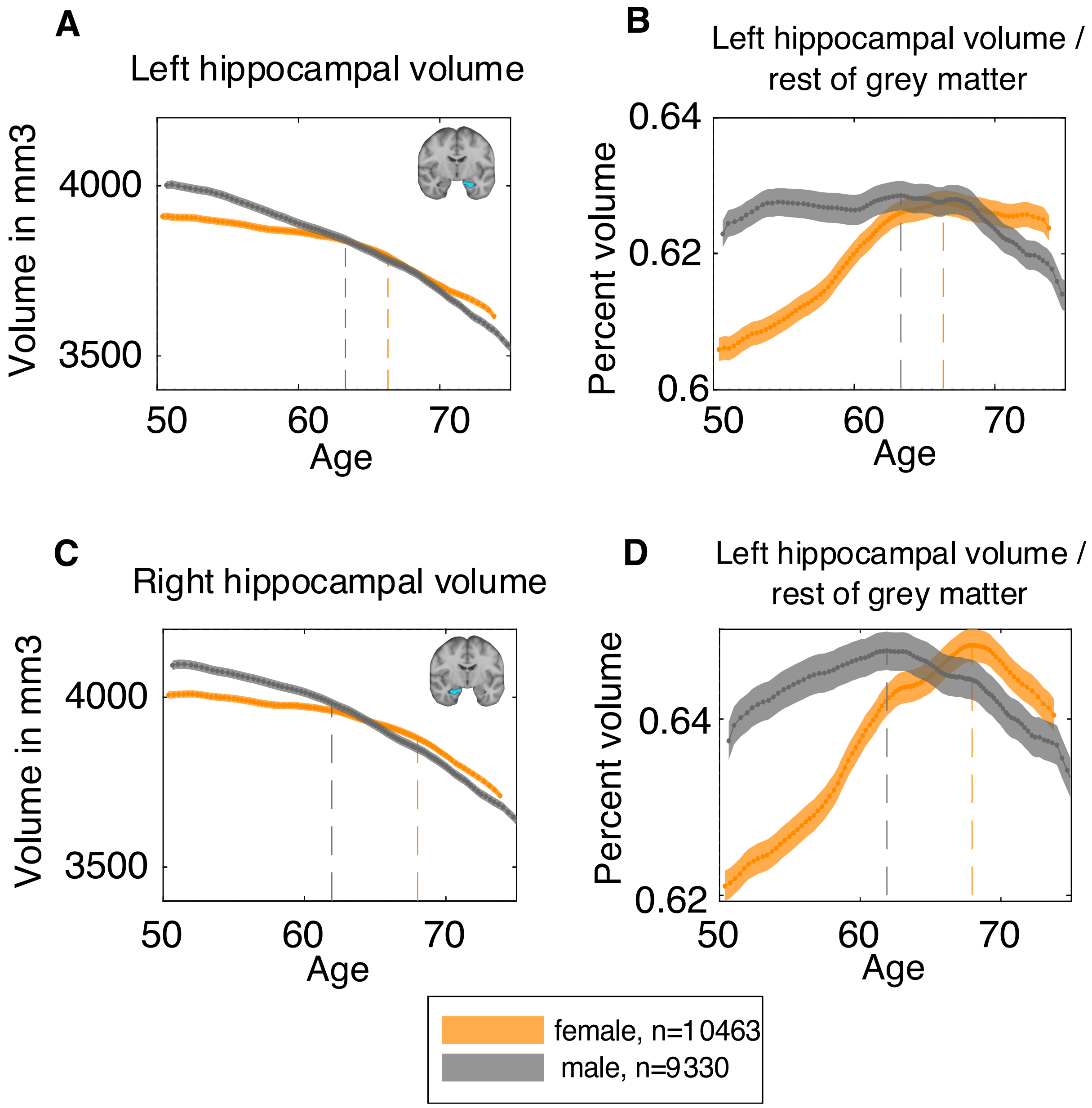


**Suppl. Figure S13**: *Sliding-window curves of left and right hippocampus across age*

**A**. Mean left hippocampal volume including standard errors as a function of age, corrected for head size. **B**. Mean left hippocampal volume to rest of grey matter ratio including standard errors as a function of age. **C**. Mean right hippocampal volume including standard errors as a function of age, corrected for head size. **D**. Mean right hippocampal volume to rest of grey matter ratio including standard errors as a function of age.

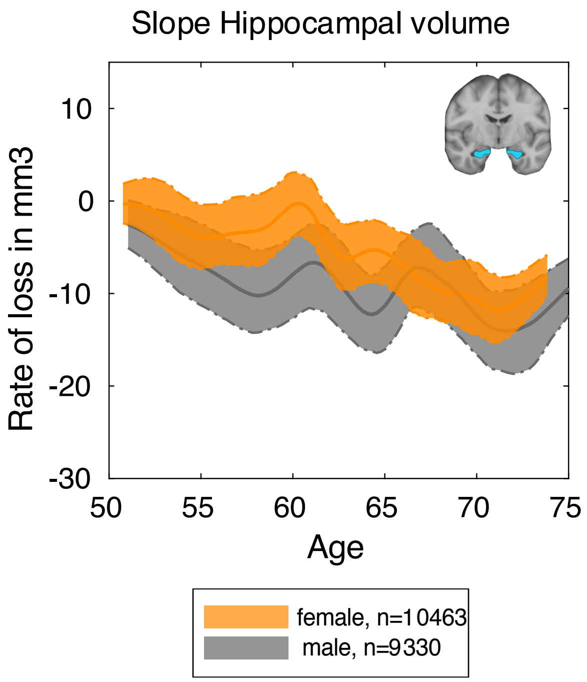


**Suppl. Figure S14**: Mean slope of bilateral hippocampal volume as a function of age, including 95% bootstrapped confidence intervals.

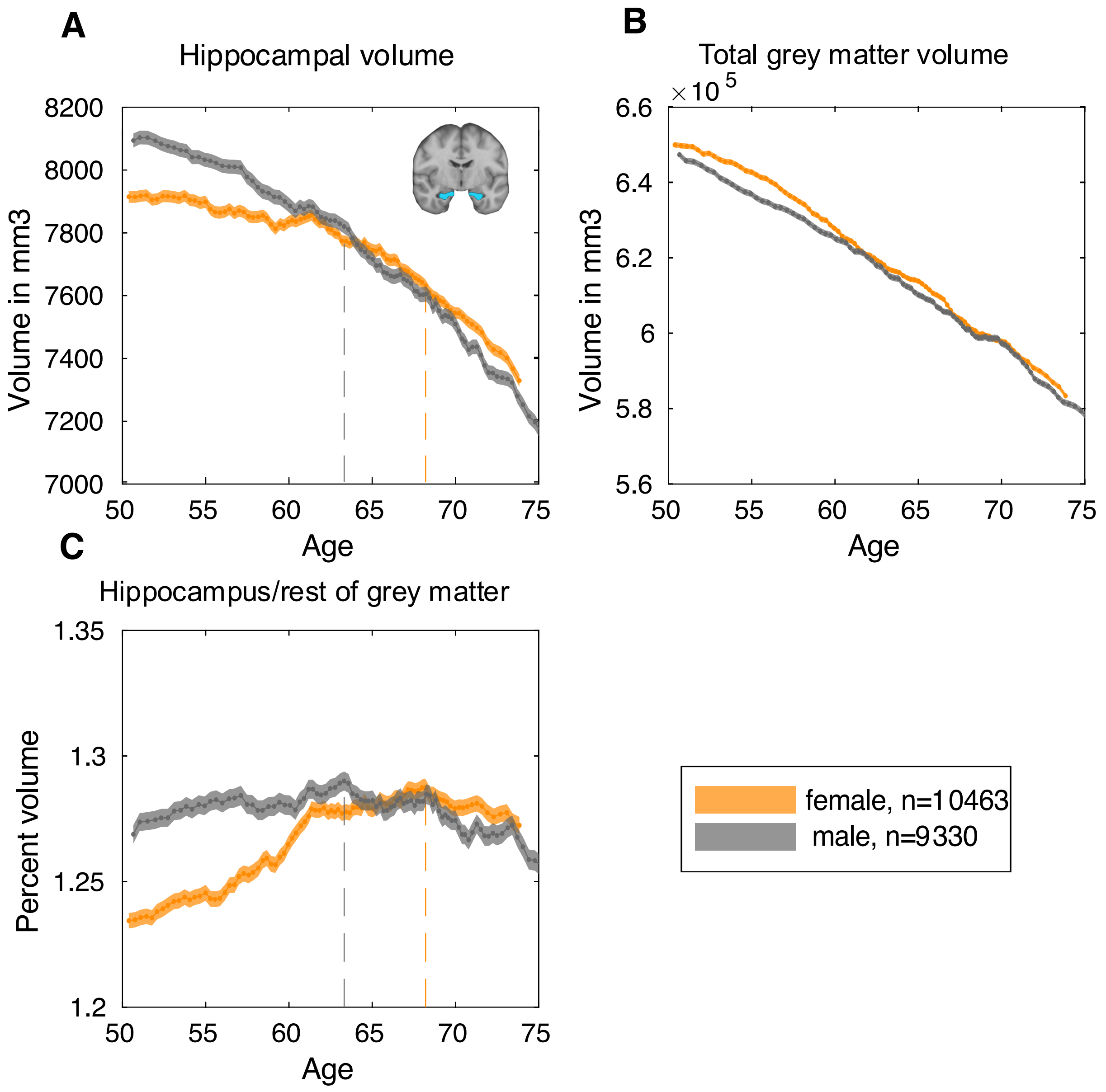


**Suppl. Figure S15:** *Sliding-window curves without smoothing*

Dashed lines indicate points of maximum ratio. **A**. Mean bilateral hippocampal volume including standard errors as a function of age, corrected for head size. **B**. Mean total grey matter volume including standard errors as a function of age, corrected for head size. **C**. Mean hippocampal volume to rest of grey matter ratio including standard errors as a function of age.

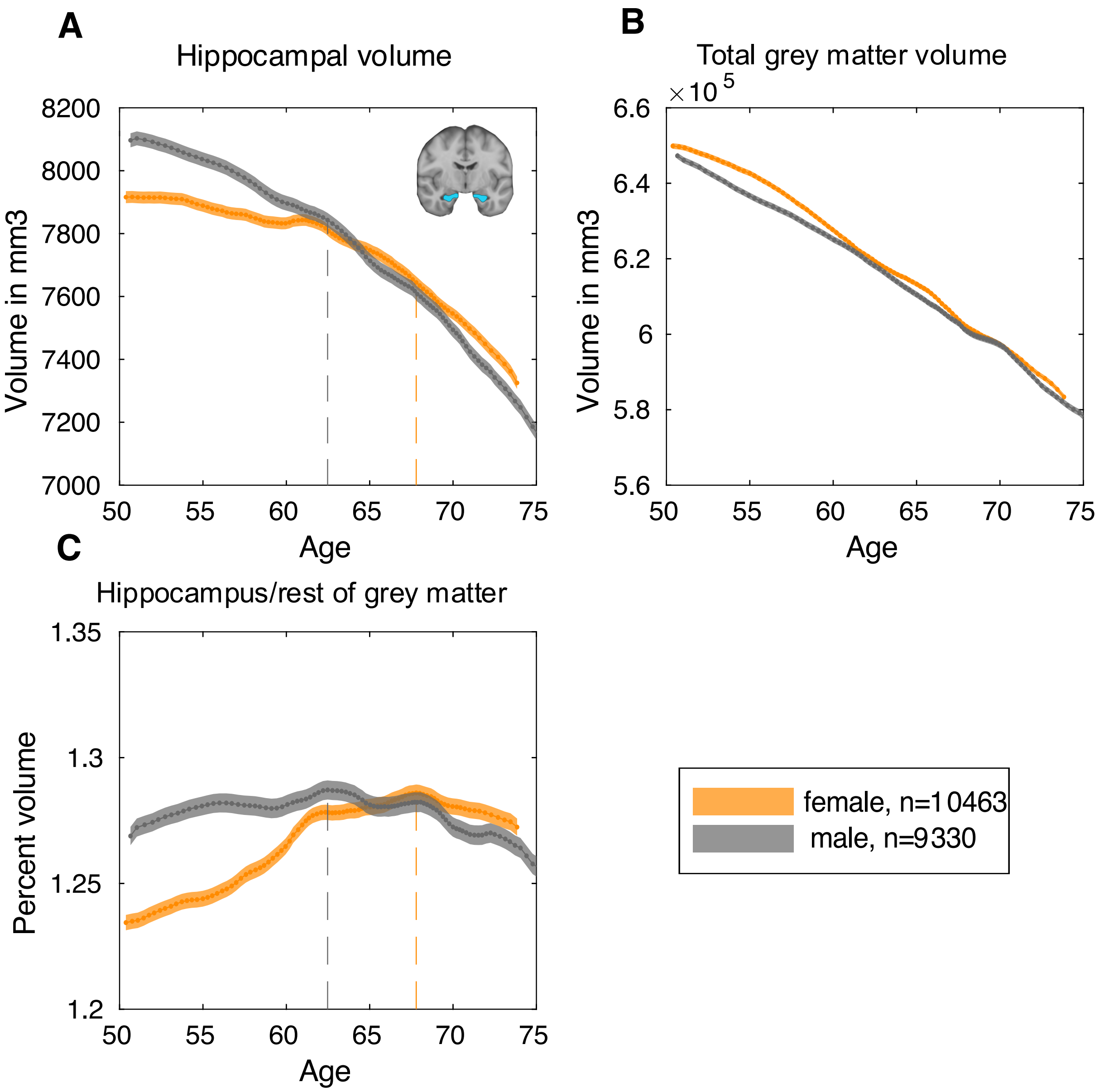


**Suppl. Figure S16:** *Sliding-window curves with smoothing kernel of 10*

Dashed lines indicate points of maximum ratio. **A**. Mean bilateral hippocampal volume including standard errors as a function of age, corrected for head size. **B**. Mean total grey matter volume including standard errors as a function of age, corrected for head size. **C**. Mean hippocampal volume to rest of grey matter ratio including standard errors as a function of age.

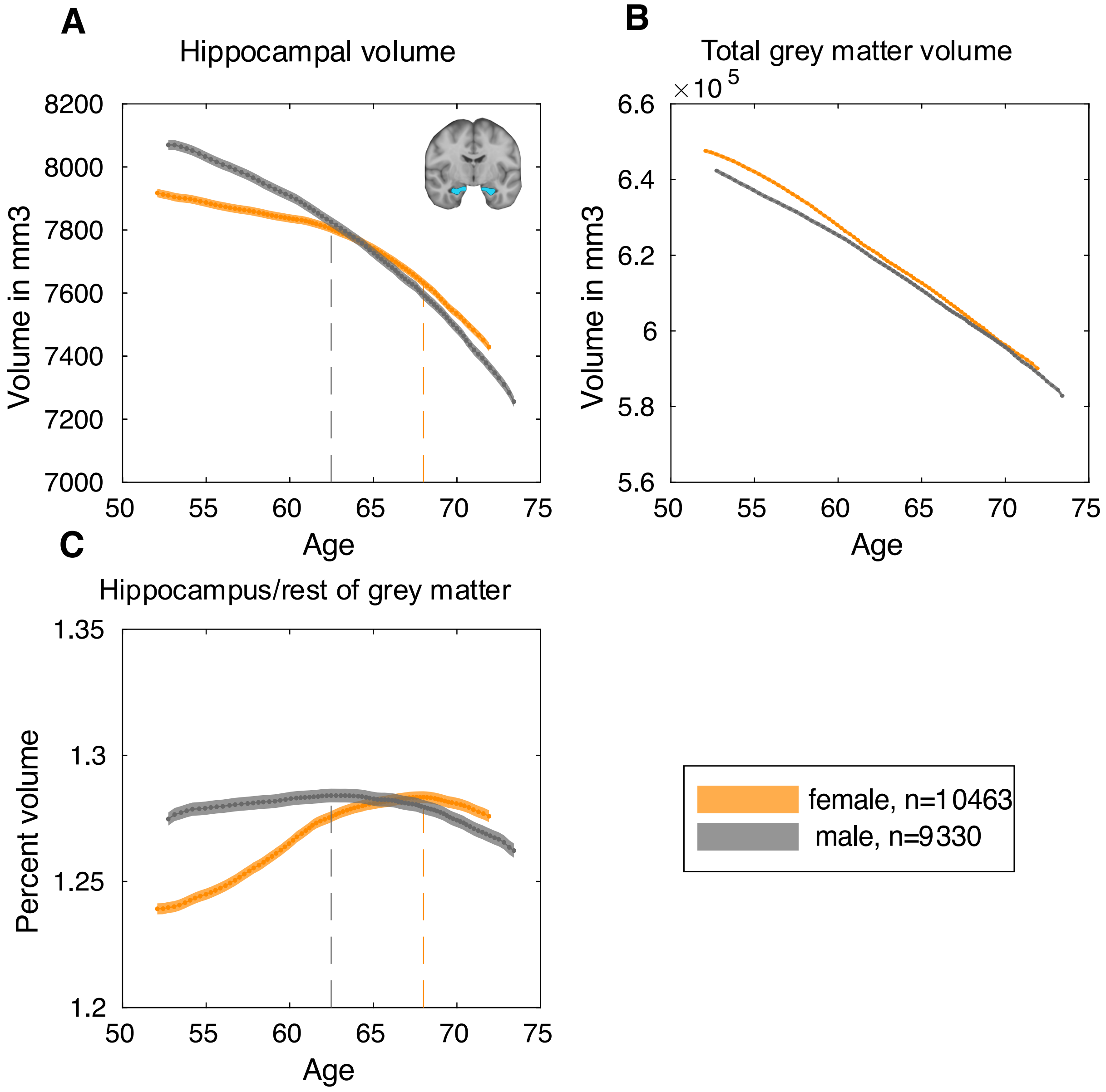


**Suppl. Figure S17:** *Sliding-window curves with 20% quantile width and smoothing kernel of 20*

Dashed lines indicate points of maximum ratio. **A**. Mean bilateral hippocampal volume including standard errors as a function of age, corrected for head size. **B**. Mean total grey matter volume including standard errors as a function of age, corrected for head size. **C**. Mean hippocampal volume to rest of grey matter ratio including standard errors as a function of age.

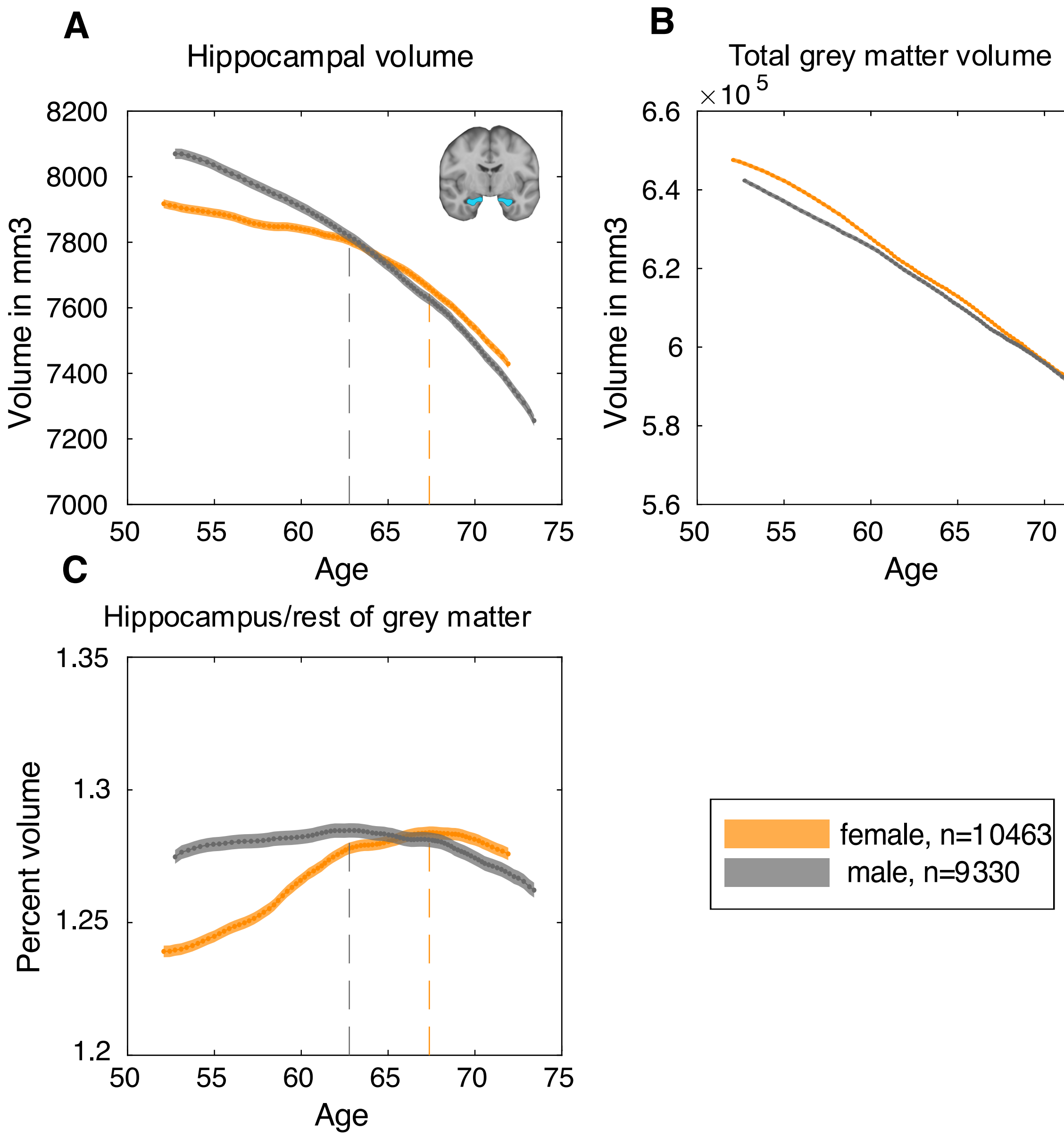


**Suppl. Figure S18:** *Sliding-window curves with quantile width 20% and smoothing kernel of 10*

Dashed lines indicate points of maximum ratio. **A**. Mean bilateral hippocampal volume including standard errors as a function of age, corrected for head size. **B**. Mean total grey matter volume including standard errors as a function of age, corrected for head size. **C**. Mean hippocampal volume to rest of grey matter ratio including standard errors as a function of age.
